# Effects of NMDA antagonists on social behaviour: a systematic review and meta-analysis of preclinical studies

**DOI:** 10.64898/2026.05.13.724847

**Authors:** Matheus Gallas-Lopes, Mariana B. Abreu, Michael Andrades, Bruno D. Arbo, Leonardo M. Bastos, Tainah C. Caetano, Daniela V. Müller, Amanda Patelli-Alves, Daniel A. Rosa, Dirson J. Stein, Ana P. Herrmann

## Abstract

Social withdrawal is a key component of the negative symptom domain of schizophrenia, and pharmacological blockade of the *N*-methyl-D-aspartate receptor (NMDAR) is widely used to model schizophrenia-relevant phenotypes in animals. However, findings on social behaviour are inconsistent across paradigms and laboratories. We therefore conducted a systematic review and meta-analysis to synthesise the effects of dizocilpine, ketamine, and phencyclidine on social interaction and social preference, to evaluate whether clinically approved antipsychotics modify these outcomes, and to examine locomotor activity measured within the same social tests to aid interpretation. We searched Embase, PubMed and Web of Science without language or date restrictions. Controlled *in vivo* studies in laboratory animals administering an eligible NMDAR antagonist and reporting social interaction and/or social preference outcomes were included. Two reviewers independently screened records, extracted data and assessed risk of bias. Effect sizes were computed as standardised mean differences and synthesised using correlated multilevel random-effects models with cluster-robust variance estimation. In total, 264 studies met the inclusion criteria. Overall, NMDAR antagonists were associated with reduced social interaction and reduced social preference relative to controls, although the social preference literature appeared vulnerable to small-study effects and imprecision. Locomotor activity measured during social interaction tests tended to be higher following NMDAR antagonists, whereas during social preference no consistent overall change was observed. In animals exposed to NMDAR antagonists, antipsychotics increased social behaviour, but these changes commonly co-occurred with reduced locomotion during social interaction tests, suggesting that improvements in social measures may partly reflect altered behavioural competition and time allocation rather than selective restoration of social functioning. Taken together, the evidence supports an overall link between NMDAR antagonism and reduced social behaviour, but the strength and interpretability of this signal depend on the paradigm and are constrained by heterogeneity and limitations in reporting.

## I. INTRODUCTION

Social withdrawal and impaired social functioning are core components of the negative symptom domain of schizophrenia and are closely associated with long-term functional disability and reduced quality of life (Galderisi *et al*., 2018; Harvey, Strassnig & Silberstein, 2019; Correll & Schooler, 2020; Mosolov & Yaltonskaya, 2022). In contrast to positive symptoms, which are responsive to antipsychotic treatment in most patients, negative symptoms demonstrate limited and inconsistent improvement with currently available pharmacotherapies, with reported benefits often attributable to changes in secondary negative symptoms (Fusar-Poli *et al*., 2015; Leucht *et al*., 2017; Krause *et al*., 2018; Correll & Schooler, 2020; Sabe *et al*., 2021). Social dysfunction, therefore, remains a persistent clinical problem and an important target for translational research (Millan *et al*., 2014; Spark *et al*., 2022; Howes, Fusar-Poli & Osugo, 2023).

Among the neurobiological hypotheses of schizophrenia, *N*-methyl-D-aspartate receptor (NMDAR) hypofunction has received substantial empirical support from convergent *post-mortem* findings (Vrajová *et al*., 2010; Weickert *et al*., 2013; Catts *et al*., 2016) and pharmacological evidence in humans (Allen & Young, 1978; Newcomer *et al*., 1999; Adler *et al*., 1999; Lahti *et al*., 2001; Pomarol-Clotet *et al*., 2006). Consistent with this framework, NMDAR antagonist-based animal models using non-competitive antagonists such as dizocilpine (MK-801), ketamine, and phencyclidine (PCP) are well established in preclinical research, being employed to elicit behavioural and neurochemical alterations related to the positive, cognitive, and negative symptoms of schizophrenia (Neill *et al*., 2010; Frohlich & Van Horn, 2014; Miyamoto & Nitta, 2014; Lee & Zhou, 2019; Benvenutti *et al*., 2022; Li *et al*., 2024).

Although social behaviour has been extensively examined in these models, reported findings remain inconsistent across the literature, with studies reporting null effects (Becker & Grecksch, 2004; Hanks *et al*., 2013) and even unexpected increases in sociability (Dunn, Corbett & Fielding, 1989; Jenkins *et al*., 2008; Rawat *et al*., 2022). The latter has also been observed in our laboratory, where acute MK-801 administration increased social behaviour in adult C57BL/6 mice (Giongo *et al*., 2023). Social outcomes are measured using heterogeneous paradigms, most commonly free social interaction and social preference tests (Toth & Neumann, 2013; Wilson & Koenig, 2014).

Social interaction paradigms quantify direct reciprocal behaviours between freely moving conspecifics (File & Hyde, 1978), whereas social preference paradigms measure approach or proximity to a constrained social stimulus (Landauer & Balster, 1982; Wee *et al*., 1995; Haller & Bakos, 2002; Nadler *et al*., 2004), also capturing motivational or exploratory components besides interaction *per se*. Furthermore, substantial variation exists in apparatus design, outcome definitions, species, developmental stage, dosing schedules, and administration routes (Neill *et al*., 2010; Benvenutti *et al*., 2022). As a result, it remains unclear to what extent the observed differences reflect biological effects of NMDAR blockade rather than methodological inconsistencies.

Interpretation is further complicated by two recurring issues. First, antipsychotics are routinely used in animal studies to test the pharmacological sensitivity of NMDAR antagonist–induced phenotypes (Spark *et al*., 2022), yet clinically available antipsychotics provide little to no benefit for primary negative symptoms in patients and have a limited impact on their social functioning (Correll & Schooler, 2020; Howes *et al*., 2023). For this reason, the apparent normalisation of social alterations in animals cannot, on its own, be taken as evidence of translational relevance (Hagan & Jones, 2005; Large, 2007; Wilson & Koenig, 2014). Therefore, it remains crucial to determine whether the reported antipsychotic effects on social outcomes reflect specific modulation of social behaviour or instead arise from non-specific changes in arousal, anxiety, or motor output, and to assess how robust these effects are across species, paradigms, and laboratories. Second, NMDAR antagonists are also known to cause hyperlocomotion (Lee & Zhou, 2019; Benvenutti *et al*., 2022). Locomotor changes can directly influence measures of both reciprocal interaction and proximity-based social behaviour (Yang, Silverman & Crawley, 2011; Mitchell *et al*., 2020), separating social effects from non-specific motor effects is thus not always straightforward.

Beyond heterogeneity in experimental design and insufficient reporting of key methodological practices, this area of research may also be affected by inadequate handling of non-independent observations (Sena *et al*., 2010; Landis *et al*., 2012; Hirst *et al*., 2014; Macleod *et al*., 2015; Wilson *et al*., 2023; Townsend *et al*., 2025). In social behaviour experiments, non-independence can arise when multiple measurements are obtained from the same animal, when interacting animals within a dyad are treated as independent experimental units, or when outcomes are clustered within cages or litters. Failure to account for such dependencies can artificially increase precision and inflate apparent effect sizes (Lazic, Clarke-Williams & Munafò, 2018; Lazic *et al*., 2020). These issues extend beyond social paradigms and reflect broader challenges in preclinical research, limiting the interpretability and reproducibility of individual studies (Wilson *et al*., 2023; Eleftheriou *et al*., 2025).

Although the existing literature suggests that NMDAR antagonists induce social behaviour deficits, the magnitude, consistency, and moderators of these effects have not been established. A quantitative synthesis is therefore required to integrate the available evidence, characterise heterogeneity, evaluate potential sources of bias, and assess the extent to which experimental factors and pharmacological interventions modulate reported effects (Bahadoran *et al*., 2020; Ineichen *et al*., 2024b). This systematic review and meta-analysis addresses the following question: what are the effects of NMDAR antagonists on social behaviour in laboratory animals? Secondary objectives are to evaluate the effects of clinically approved antipsychotics on social outcomes in NMDAR antagonist–exposed animals and to assess locomotor activity as a control outcome to aid interpretation of social measures.

## II. METHODS

### (1) Protocol

The protocol for this systematic review and meta-analysis was registered in the International Prospective Register of Systematic Reviews (PROSPERO, registration number: CRD42023402129) prior to study screening and data extraction (Gallas-Lopes *et al*., 2024). The preregistered protocol specified the research questions, eligibility criteria, search strategy, and planned analyses.

The primary objective of the review was to evaluate the effects of NMDAR antagonists (MK-801, ketamine, or PCP) on social behaviour in laboratory animals, with social behaviour operationalised as social interaction or social preference outcomes. Secondary objectives were to assess the effects of clinically approved antipsychotics on social behaviour in animals exposed to NMDAR antagonists, and to evaluate locomotor activity in both comparisons (NMDAR antagonist vs control, and NMDAR antagonist plus antipsychotic vs NMDAR antagonist alone) as a control outcome to determine whether observed social behaviour changes could be confounded by alterations in general activity levels.

This review adheres to the Preferred Reporting Items for Systematic Reviews and Meta-Analyses (PRISMA) guidelines (Page *et al*., 2021a, 2021b). All data and materials associated with this study, including extracted datasets and analysis code, are publicly available via the Open Science Framework (OSF) at osf.io/98fau (Gallas-Lopes *et al*., 2023a) and GitHub at github.com/matheusglls/NMDA-sb (Gallas-Lopes, 2026).

### (2) Study selection

Systematic searches were conducted in Embase, PubMed, and Web of Science on February 25^th^, 2023. The search strategy was designed to capture studies evaluating the effects of NMDAR antagonists on social behaviour in laboratory animals and combined controlled vocabulary and free-text terms relating to the population (laboratory animals), intervention (NMDAR antagonists), and outcomes (social interaction and social preference). For the population component, validated SYRCLE animal filters were used to ensure broad coverage of laboratory animal studies across species (van der Mierden *et al*., 2022). No restrictions were applied on language or publication date. Full database-specific search strategies were preregistered and are available at https://osf.io/5k2br (Gallas-Lopes *et al*., 2023b).

Records retrieved from the database searches were exported and imported to Rayyan (Ouzzani *et al*., 2016), where duplicate entries were identified and removed using a combination of automated detection and manual verification. Study selection was carried out in two phases: an initial screening based on titles and abstracts, followed by a full-text eligibility assessment. In both phases, each record was evaluated independently by two reviewers, and decisions were made blinded to the other reviewer’s judgment.

During the title and abstract screening stage, records were excluded only when they clearly failed to meet eligibility criteria related to study design, population, or intervention. Specifically, studies were excluded if they were not original controlled experiments, did not involve live laboratory animals, or did not include administration of eligible NMDAR antagonists (MK-801, ketamine, or PCP). Records for which eligibility could not be confidently determined were retained for full-text assessment. In the second stage, full texts of potentially eligible articles were assessed in detail using the same criteria applied in the first stage, with the addition of criteria related to the presence of an appropriate control group and the assessment of relevant social behaviour outcomes (social interaction and/or social preference). Any disagreements were resolved through discussion, and when consensus was not achieved, a third reviewer was consulted.

All screening decisions followed a standardised reviewer guide detailing operational definitions of inclusion and exclusion criteria and specifying hierarchical exclusion rules available at osf.io/98fau/files/3pm2n. This guide was developed before screening and applied throughout both screening phases. All screening decisions were managed and archived within Rayyan.

### (3) Data collection

Data extraction was performed in two sequential stages: an initial qualitative extraction to characterise study and experimental features, followed by quantitative extraction of outcome data for meta-analysis. Qualitative and quantitative extractions were conducted following predefined guides, available at osf.io/98fau/files/k928m and osf.io/98fau/files/bhdp3, respectively. Both stages were conducted independently by two reviewers using standardised extraction forms developed before data collection. When discrepancies occurred or when differences between extracted quantitative values exceeded 10%, reconciliation was performed by a third reviewer who re-extracted the discrepant information to establish final values.

In the qualitative extraction stage, descriptive information was collected for each experiment, including publication details, animal characteristics (species, strain, sex, age, or developmental stage at both induction and outcome assessment), and intervention parameters (NMDAR antagonist used, dose or concentration, route of administration, frequency and duration of exposure, and interval between drug administration and behavioural testing). Characteristics of the behavioural assessment were also recorded, including the type of test (social interaction or social preference) and the specific outcome measure.

Following qualitative characterisation, quantitative extraction was performed for all eligible social behaviour outcomes. For each comparison, group-level summary statistics were extracted, including sample size, mean values, and measures of variability (standard deviation or standard error of the mean). When data were reported in text or tables, exact values were preferentially extracted from those sources. When numerical values were not provided, data were estimated from figures using WebPlotDigitizer (https://automeris.io/). If it was unclear whether error bars represented standard deviation or standard error, values were conservatively treated as standard error and later converted to standard deviation for effect size estimation. When the sample size was reported as a range, the lowest value was used for effect size estimation, whereas the highest value was used for calculating standard deviation when it had to be derived from the standard error of the mean. When pseudoreplication was suspected, sample sizes were adjusted to reflect the appropriate biological unit. For example, when social behaviour was quantified separately for each animal within interacting pairs, the effective sample size was divided by two to account for non-independence within dyads.

Experiments conducted in distinct sets of animals within the same study were entered as separate effect sizes, while dependence among effect sizes from the same study was accounted for using multilevel meta-analytic models. Studies employing cross-over designs or reusing the same animals across multiple experimental conditions were excluded when it was impossible to isolate results obtained exclusively from naive animals. When multiple experiments were conducted sequentially in the same animals, only data from the first behavioural assessment were retained. Locomotor activity data were extracted only when locomotion was measured in the same animals and within the same experimental context as the social behaviour assessment. When outcomes were assessed at multiple time points within the same experiment, only the earliest post-intervention time point was considered. Outcomes exclusively assessing social novelty preference were not considered eligible. When essential quantitative information, such as sample size or variability measures, was missing or could not be reliably obtained, corresponding authors were contacted by email on up to two occasions. Studies were excluded when required data could not be retrieved. Studies reporting outcomes exclusively as medians with interquartile ranges were excluded from the quantitative synthesis when raw data were not available to calculate means and standard deviations, and when additional data were not provided by the authors.

### (4) Risk of bias

Risk of bias was assessed using SYRCLE’s risk of bias tool for animal studies (Hooijmans *et al*., 2014b), with additional domains addressing pseudoreplication and procedural equivalence. Pseudoreplication refers to the inappropriate treatment of non-independent observations (e.g., multiple measurements from the same animal or cage) as independent experimental units, which can inflate precision and bias effect estimates (Lazic *et al*., 2018). Procedural equivalence assesses whether experimental groups were handled identically aside from the intervention (e.g., same housing conditions, handling, route of administration, timing, and outcome assessment procedures), minimising performance and detection bias. Due to pervasive poor reporting in animal studies, the allocation concealment domain was omitted, as it cannot be objectively evaluated in most included studies (Hirst *et al*., 2014). A detailed assessment guide with operational definitions and decision rules for each domain is publicly available at osf.io/98fau/files/5gkct.

Only experiments and behavioural outcomes relevant to this review were considered when assigning risk of bias judgments. When a study included more than one eligible experiment or behavioural test, judgments were made for each relevant experiment and then summarised at the study level. This summary followed a conservative approach: if any relevant experiment within a study was judged as high risk in a given domain, the study was classified as high risk for that domain. Domains included sequence generation, baseline characteristics, random housing, blinding during the experiment, random outcome assessment, blinding during outcome assessment, incomplete outcome data, selective outcome reporting, pseudoreplication, and procedural equivalence. For each domain, risk of bias was rated as low, unclear, or high. An overall study-level classification was then assigned and dichotomised as low or high, with studies rated as high overall risk of bias when at least one domain was judged as high risk. All assessments were performed independently by two reviewers, and discrepancies were resolved by discussion, with conciliation by a third reviewer when necessary.

Inter-rater reliability was quantified based on initial independent assessments using observed agreement (i.e., the proportion of identical ratings between assessors), Cohen’s kappa (Cohen, 1960), and prevalence-adjusted bias-adjusted kappa (PABAK) (Byrt, Bishop & Carlin, 1993), calculated overall and by domain. Risk of bias results were summarised descriptively across domains and studies using summary bar plots and traffic-light plots. All analyses and visualisations were conducted in R (version 4.3.1). The full analysis scripts, package versions, and raw data are publicly available via OSF (https://osf.io/98fau) and GitHub (github.com/matheusglls/NMDA-sb/tree/main/risk_of_bias).

### (5) Outcome selection and effect size estimation

When more than one outcome assessing the same behavioural domain (social preference, social interaction, or locomotor activity) was extracted for the same group of animals within an experiment, only one outcome was selected for inclusion in the meta-analysis. This approach was adopted to avoid multiple dependent effect sizes arising from the same animals. Outcome selection followed a predefined hierarchy based on the frequency in which specific outcome types were reported across studies and on how directly each measure captured the behavioural construct of interest. Outcomes with different names but assessing the same underlying construct were grouped and ranked accordingly.

For social preference and social interaction, time-based measures reflecting the absolute duration of social behaviour were prioritised, as these were the most frequently reported and provided the most direct quantification of social behaviour. When time-based measures were not available, we selected the available measure that most directly represented the same construct, prioritising proportion- or index-based measures of social behaviour. Latency- or distance-based outcomes (e.g., latency to first contact, distance to the stimulus) were selected only when no other measures were reported.

For locomotor activity, absolute measures of locomotor activity based on total distance or movement trajectories were prioritised. When these were not available, count-based measures of activity, such as line crossings, sector crossings, beam breaks, or locomotor/motor activity counts, were selected, followed by time-based measures, such as activity time or duration of locomotion. Relative, normalised, or derived locomotor measures were included when no absolute locomotor outcomes were reported.

Effect sizes were calculated as standardised mean differences using the Hedges’ g method, with corresponding sampling variances estimated using the *escalc* function of the *metafor* package in R (version 4.3.1) (Viechtbauer, 2010). To ensure consistent directionality across outcomes, effect sizes were reversed when lower values indicated higher levels of social behaviour, such as latency to first social contact or measures based on inter-individual distance.

The distribution of effect sizes included in the meta-analyses was summarised visually to characterise the structure of the available evidence. Alluvial plots were used to illustrate how effect sizes were distributed across key study characteristics and were constructed using the *ggalluvial* package (Brunson & Read, 2023). Evidence maps were constructed to summarise the number of effect sizes and their mean magnitude across combinations of experimental factors, providing an overview of data density and coverage.

### (6) Data synthesis

Meta-analyses were conducted separately for two conceptually distinct domains of social behaviour: social interaction and social preference. Social interaction paradigms were defined as those allowing direct physical interaction between two or more conspecifics. In contrast, social preference paradigms assessed interest in or proximity to a conspecific whose movement or physical contact was restricted (e.g., wire cages or enclosed compartments). Locomotor activity outcomes assessed during social behaviour tests were analysed in parallel with each social domain.

Based on these distinctions and the experimental contrasts of interest, seven independent meta-analyses were performed:

1. Effects of NMDAR antagonists on social interaction;
2. Effects of NMDAR antagonists on locomotor activity measured during social interaction paradigms;
3. Effects of NMDAR antagonists on social preference;
4. Effects of NMDAR antagonists on locomotor activity measured during social preference paradigms;
5. Effects of antipsychotics on NMDAR antagonist–induced alterations in social interaction;
6. Effects of antipsychotics on locomotor activity measured during social interaction paradigms in animals exposed to NMDAR antagonists;
7. Effects of antipsychotics on NMDAR antagonist–induced alterations in social preference.

Effects of antipsychotics on locomotor activity measured during social preference paradigms in NMDAR antagonist–exposed animals were not meta-analysed, as no eligible comparisons reported this outcome within the same animals and behavioural context.

Effect sizes were synthesised using correlated multilevel random-effects models, in which effect sizes were nested within experiments and experiments were nested within studies, to properly account for dependencies. Random intercepts for both study and experiment were included to capture between-study and within-study variability. Models were fitted using restricted maximum likelihood estimation. In addition, cluster-robust variance estimation at the study level was applied to all models using the *clubSandwich* package (Pustejovsky, Pekofsky & Zhang, 2025), providing additional protection against residual dependence.

To account for shared control groups and resulting dependency between effect sizes derived from the same experiment, correlations were modelled using a variance–covariance matrix constructed with the *vcalc* function. A within-experiment correlation coefficient (ρ) of 0.5 was assumed.

Heterogeneity was examined to evaluate how much effect sizes varied beyond sampling error. Cochran’s Q statistic was used to test whether heterogeneity was present. The magnitude of heterogeneity was quantified using multilevel I² estimates (Nakagawa *et al*., 2023) and variance components (σ²) from the random-effects models, which allowed heterogeneity to be assessed separately at the study and experiment levels.

Pooled effects were summarised graphically using orchard plots constructed with the *orchaRd* package (Nakagawa *et al*., 2023), which display study-level effect estimates alongside the pooled effect while accounting for multilevel heterogeneity and within-study dependency. To complement confidence intervals around pooled estimates, prediction intervals were calculated to indicate the expected range of true effects that might be observed in future studies, incorporating both sampling error and between-study heterogeneity.

Small-study effects, which may reflect publication bias, were examined using funnel plots and regression-based methods. Funnel plots were constructed using standard error and the inverse square root of the total sample size to assess asymmetry (Egger *et al*., 1997; Sterne *et al*., 2011; Zwetsloot *et al*., 2017). Precision-effect test (PET) and precision-effect estimate with standard error (PEESE) models were fitted by regressing effect sizes on standard error and sampling variance, respectively, within the multilevel framework. Following recommended practice, when the PET indicated evidence of small-study effects (i.e., a significant slope), the PEESE estimate was interpreted as the primary bias-adjusted effect size; otherwise, the PET estimate was retained (Stanley, 2008; Stanley & Doucouliagos, 2014). As an additional sensitivity analysis, we performed a regression using a standard error adjusted for effective sample size. This approach reduces the artifactual correlation between standardized mean differences and their standard errors, providing a complementary assessment of small-study effects. Time-lag bias was assessed using meta-regression with publication year centred around its mean and included as a continuous moderator (Yang *et al*., 2023).

Meta-regression analyses were conducted within the multilevel framework to examine whether the effects of NMDAR antagonists on social interaction and social preference varied across key experimental and biological characteristics and to assess whether these variables explained part of the between-study heterogeneity. A priori planned categorical moderators included NMDAR antagonist, species, developmental stage at induction, route of administration, dosing schedule, sex, and, when applicable, antipsychotic treatment. All moderator models were fitted; however, only moderators for which each predictor level was informed by at least five independent studies were considered sufficiently powered and therefore reported and discussed in the main text. For the remaining moderators, sparse data within individual levels limited the precision and interpretability of estimates.

We also performed meta-regressions to examine dose–response patterns using cumulative NMDAR antagonist exposure as a continuous predictor. This analysis was restricted to studies conducted in rats and mice, where reporting of dosing parameters was sufficiently consistent to derive cumulative exposure estimates. When data were available, cumulative exposure was calculated as the product of the administered dose (mg/kg), intervention frequency (number of administrations per day), and duration of exposure (days). This allowed single-administration and repeated-dosing schedules to be expressed on a common exposure scale. To address potential non-linearity and reduce skewness, cumulative exposure was log₁₀-transformed before analysis. The meta-regression was fitted within the same correlated multilevel random-effects framework, with robust variance estimation.

Multivariate meta-regression models were used to evaluate whether variation in effect sizes could be explained by multiple experimental characteristics simultaneously. Moderators included species, type of NMDAR antagonist, dosing schedule, and developmental stage at induction. Including these predictors jointly allowed the estimation of the independent association of each characteristic while accounting for potential confounding due to imbalance in study designs.

For meta-regression and multivariable models, marginal and conditional R² values were calculated to quantify the proportion of heterogeneity explained by the moderators alone (marginal R²) and by the full model including both moderators and random effects (conditional R²) (Nakagawa & Schielzeth, 2013).

The full analytical pipeline was applied to the primary meta-analyses evaluating the effects of NMDAR antagonists on social interaction and social preference, which constituted the main confirmatory analyses of this review. For these outcomes, analyses included multilevel random-effects models, categorical moderator analyses, multivariable meta-regression, cumulative exposure meta-regression, time-lag analyses, assessment of small-study effects, and sensitivity analyses. For locomotor activity outcomes and antipsychotic comparisons, analyses were restricted to the core meta-analytic model and selected sensitivity analyses because of more limited and heterogeneous data structures. The primary meta-analyses are therefore emphasised in the main text.

All analyses were conducted in R (version 4.3.1). The full analysis scripts, package versions, and raw data are publicly available via OSF (https://osf.io/98fau) and GitHub (github.com/matheusglls/NMDA-sb/tree/main/meta_analysis).

### (7) Sensitivity analyses

Sensitivity analyses were conducted using the same correlated multilevel random-effects models with robust variance estimation applied in the primary analyses. First, to assess the robustness of pooled estimates to the assumed within-experiment correlation used in the variance–covariance matrix, models were re-estimated using alternative values of ρ across a plausible range (0.0, 0.3, 0.5, and 0.8). Second, to assess the influence of individual studies, leave-one-study-out analyses were performed by sequentially excluding each study and re-estimating the pooled effect, allowing evaluation of whether the overall results were disproportionately driven by any single study. Third, analyses were repeated after excluding studies classified as having high overall risk of bias, to examine whether conclusions were sensitive to the inclusion of studies with greater methodological limitations. Consistency between the primary and sensitivity analyses was interpreted as evidence of robustness.

### (8) Deviations from protocol

Some qualitative variables were extracted as prespecified, yet were not analysed because reporting was unclear or incomplete in most studies. Locomotor activity was initially planned for extraction whenever reported; however, we ultimately extracted it only when it was measured during the social test and in the same animals, to better evaluate whether locomotor changes could confound social outcomes. Locomotion measured in separate apparatuses or in different animals was therefore not included in our analyses. In addition, the data synthesis approach and risk of bias analyses were adapted to better accommodate dependent effect sizes and the multilevel structure of the included evidence. Although several subgroup and moderator analyses were prespecified, many predictor levels were informed by too few studies to allow reliable estimation. Finally, we conducted a non-prespecified dose-response meta-regression in rats and mice using log-transformed cumulative NMDAR antagonist exposure.

## III. RESULTS

Visualisations of the primary and secondary analyses are provided at matheusglls.github.io/NMDA-sb.

### (1) Study selection

A total of 2,261 records were identified through database searches, including Embase (*n* = 913), PubMed (*n* = 708), and Web of Science (*n* = 640). After the removal of 941 duplicate records, 1,320 records remained and were screened based on titles and abstracts. Of these, 401 records were excluded, primarily due to study design (*n* = 204), intervention (*n* = 183), and population (*n* = 14) not meeting the eligibility criteria. Consequently, 919 reports were sought for retrieval, of which 8 reports could not be retrieved. The remaining 911 reports underwent full-text assessment for eligibility. At this stage, 647 reports were excluded for not meeting the following criteria: study design (*n* = 147), population (*n* = 11), intervention (*n* = 172), comparison (*n* = 12), outcome of interest (*n* = 288), and insufficient or missing data (*n* = 17). Ultimately, 264 studies met the inclusion criteria and were included in the quantitative synthesis (Figure 1), including studies in rats (*Rattus* spp.), mice (*Mus* spp.), and zebrafish (*Danio rerio*). The complete list of included studies is available at https://osf.io/98fau/files/5qevp.

**Figure 1.**
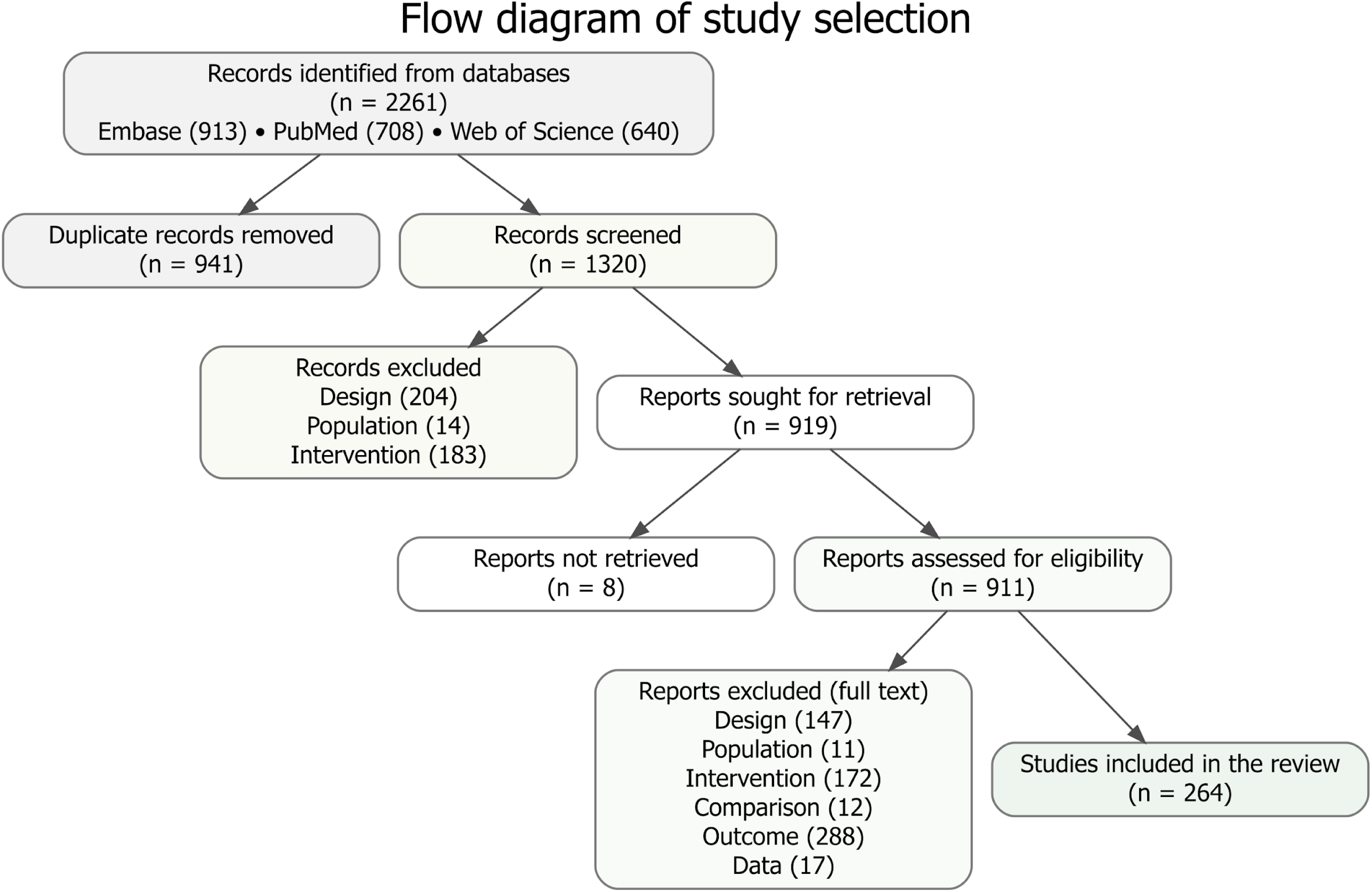
Flow diagram of the literature search and study selection process.

### (2) Risk of bias

Across domains, most studies were judged as having an unclear risk of bias, reflecting widespread under-reporting of key methodological details in the preclinical literature (Figure 2). This was particularly evident for domains related to randomisation procedures. Random sequence generation was judged as unclear in 98% of studies. Similarly, random housing was rated as unclear in 98% of studies, and random outcome assessment was unclear in 97%, indicating that these procedures were rarely reported in sufficient detail to allow confident judgment. Baseline characteristics were unclear in 61% of studies and judged as low risk in 39%.

**Figure 2.**
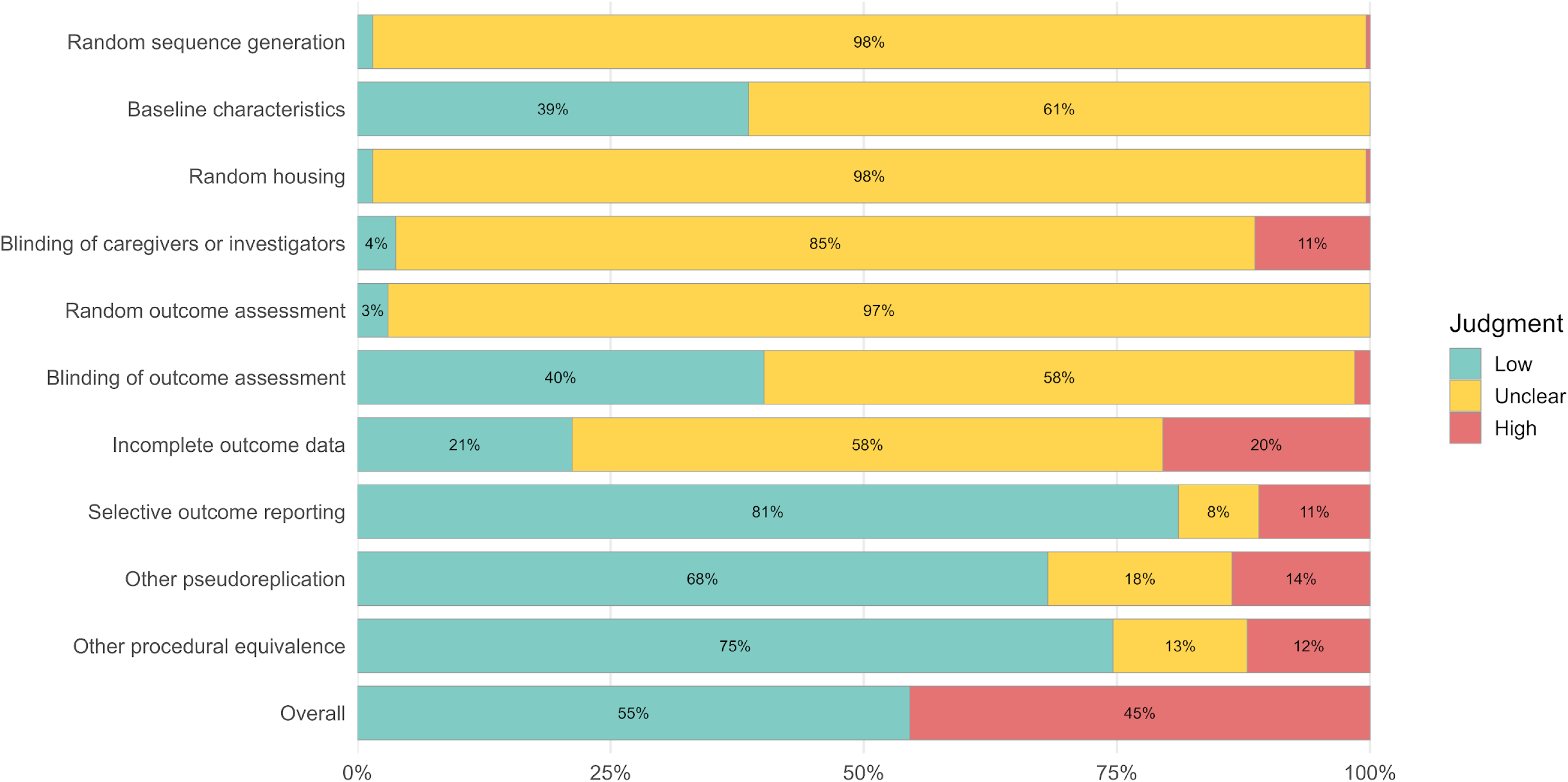
Risk of Bias summary across domains in the included studies. Bars represent the proportion of studies rated as low, unclear, or high risk of bias for each domain, based on SYRCLE’s risk of bias tool with additional domains for pseudoreplication and procedural equivalence. Domains with a higher proportion of low-risk judgments indicate stronger methodological rigour, whereas domains dominated by unclear or high risk suggest potential limitations due to insufficient reporting or methodological concerns. The “Overall” category reflects the study-level risk of bias assessment.

Blinding procedures were also inconsistently reported. Blinding of caregivers or investigators was judged as unclear in 85% of studies, low risk in 4%, and high risk in 11%. Blinding of outcome assessment showed comparatively better reporting, with 40% of studies classified as low risk, 58% as unclear, and 2% as high risk.

Domains related to data handling and reporting showed greater variability. Incomplete outcome data were judged as low risk in 21% of studies, unclear in 58%, and high risk in 20%. Selective outcome reporting was more frequently judged as low risk (81%), although 8% of studies were classified as unclear and 11% as high risk. Because preregistered protocols were generally unavailable, these judgments were based primarily on whether outcomes described in the methods were reported in the results.

For design-related domains, pseudoreplication was rated as low risk in 68% of studies, unclear in 18%, and high risk in 14%. High-risk judgments were primarily driven by an inappropriate definition of the experimental unit, especially in social interaction paradigms where individual animals within the same interacting dyad were treated as independent observations. Procedural equivalence was judged as low risk in 75% of studies, unclear in 13%, and high risk in 12%, indicating that most studies applied comparable procedures across groups, although methodological concerns remained in a minority of cases. When summarised across all domains, 55% of studies were classified as overall low risk of bias and 45% as high risk.

Inter-rater reliability for the risk of bias assessment indicated an adequate level of consistency between reviewers, with a Cohen’s kappa of 0.643 and a PABAK of 0.627. Of all ratings, 81.3% (*n* = 2,147) represented agreement between reviewers, while 18.7% (*n* = 493) reflected disagreement. Critical disagreement, defined as opposing judgments at the extremes of the scale (low vs high risk), accounted for 4.8% (*n* = 128) of all ratings.

Inter-rater reliability varied across individual risk of bias domains (Figure S1). Observed agreement was highest for sequence generation (0.981), random housing (0.981), and random outcome assessment (0.958); however, corresponding Cohen’s kappa values were moderate (0.538, 0.437, and 0.333, respectively), likely reflecting the high prevalence of “unclear” ratings in these domains. Blinding of outcome assessment showed both high observed agreement (0.883) and high agreement beyond chance (kappa 0.765). Lower levels of agreement were observed for incomplete outcome data (observed agreement 0.682, kappa 0.439), selective outcome reporting (observed agreement 0.769, kappa 0.215), pseudoreplication (observed agreement 0.557, kappa 0.245), and procedural equivalence (observed agreement 0.712, kappa 0.179), indicating greater subjectivity in judging these items.

Study- and domain-level risk of bias assessments are reported in the online resource at matheusglls.github.io/NMDA-sb/risk_of_bias/rob_and_interrater.html, together with additional analyses of inter-rater reliability.

### (3) Meta-analyses

#### (a) Effects of NMDAR antagonists on social interaction

All qualitative mappings of study characteristics and the complete results of the analyses conducted for the effects of NMDAR antagonists on social interaction are available at matheusglls.github.io/NMDA-sb/meta_analysis/nmda_antagonist_effects/social_interaction/nmda_ effects_social_interaction.html.

##### (i) Qualitative synthesis

Studies with rats contributed the greatest number of effect sizes (*k* = 341 from 123 studies), followed by mice (*k* = 141 from 57 studies), with a small number obtained from zebrafish (*k* = 9 from 6 studies). For the NMDAR antagonists, PCP accounted for the largest number of effect sizes (*k* = 227 from 78 studies), followed by MK-801 (*k* = 174 from 71 studies) and ketamine (*k* = 90 from 43 studies). The distribution of effect sizes across species and NMDAR antagonists is summarised in Figure 3A, which illustrates the relative contribution of each species and antagonist combination.

**Figure 3.**
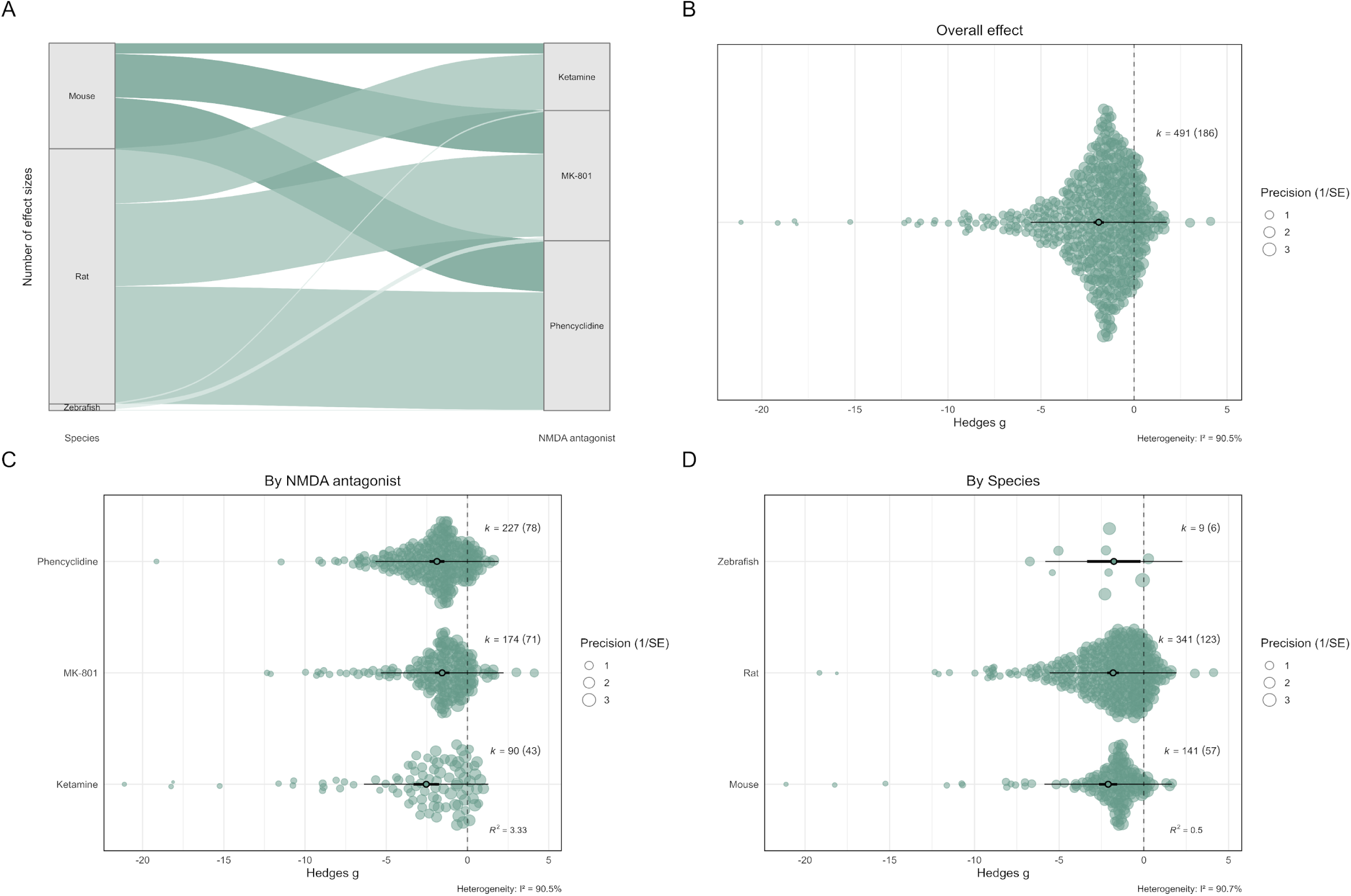
Effects of NMDA receptor antagonists on social interaction in laboratory animals. (A) Alluvial plot showing the distribution of effect sizes across species and NMDA antagonists. Flow width represents the number of effect sizes contributing to each category. (B) Orchard plot of the overall pooled effect of NMDA receptor antagonists on social interaction. (C) Orch ard plots showing effects stratified by NMDA antagonist. (D) Orchard plots showing effects stratified by species. Filled circles represent individual effect sizes scaled by study precision, and open circles indicate pooled point estimates with 95% confidence intervals (thick horizontal lines) and prediction intervals (thin horizontal lines). k denotes the number of effect sizes, with the number of independent studies shown in parentheses.

Most effect sizes were obtained following intraperitoneal administration (*k* = 240 from 112 studies), representing the dominant administration route across species and antagonists (Figure S2-3). Subcutaneous administration was also frequently used (*k* = 224 from 62 studies). Other administration routes were sparsely represented, including immersion (*k* = 7 from 4 studies), minipump delivery (*k* = 9 from 4 studies), retro-orbital injection (*k* = 1 from 1 study), combined intraperitoneal and subcutaneous administration (*k* = 1 from 1 study), and unclear administration route (k = 9 from 4 studies).

When considering the dosing schedule, effect sizes were most commonly derived from repeated administrations (*k* = 284 from 110 studies) and acute administration (*k* = 198 from 79 studies), with continuous exposure observed in only a small number of studies (*k* = 9 from 4 studies) and restricted to MK-801 and PCP experiments (Figure S4).

The developmental timing of induction varied considerably across studies. Adult and juvenile/adolescent animals accounted for the majority of effect sizes (each *k* = 123, from 63 and 44 studies, respectively), while neonatal exposure contributed relatively few (*k* = 28 from 10 studies). In addition, a large number of effect sizes were associated with studies that did not clearly report the developmental stage at induction (*k* = 217 from 70 studies) (Figure S5).

There was a marked imbalance in sex representation. Most effect sizes were derived from male animals (k = 460, 93.7%; mouse: k = 132; rat: k = 328), whereas female-only samples (k = 16, 3.3%; mouse: k = 2; rat: k = 10; zebrafish: k = 4) and mixed-sex samples (k = 11, 2.2%; mouse: k = 5; rat: k = 1; zebrafish: k = 5) were rarely used. Only a small number of effect sizes were drawn from studies in which sex was unclear (k = 4, 0.8%; mouse: k = 2; rat: k = 2) (Figure S6).

##### (ii) Quantitative synthesis

The quantitative synthesis included 491 effect sizes from 186 independent studies. Exposure to NMDAR antagonists was associated with lower social interaction relative to controls (*Hedges’ g* = −1.90, 95% CI [−2.18, −1.62], with a prediction interval of −5.56 to 1.76; Figure 3B). Heterogeneity was high (*I²_total_* = 90.5%), with substantial between-study variability (*I²_study_* = 84.3%) and a smaller contribution attributable to within-study clustering (*I²_study/exp_*= 6.2%). The test for heterogeneity was significant (*Q_490_* = 3230.49, p < 0.001), indicating considerable dispersion in effects across studies. The estimated variance components were *σ²_study_* = 3.179 and *σ²_study/exp_* = 0.233.

Visual inspection of the funnel plots (Figure S7) suggested asymmetry, consistent with the presence of publication bias. PET indicated a significant association between effect size and study precision (*slope* = −5.94, *p* < 0.001; *marginal R²* = 0.906; *conditional R²* = 1.000), providing evidence of small-study effects. PEESE also identified a significant association between effect size and sampling variance (*slope* = −1.77, *p* = 0.001). The bias-adjusted intercept from the PEESE was −0.72, 95% CI [−1.16, −0.29], indicating that adjustment for small-study effects reduced the magnitude of the effect. Residual heterogeneity remained high (*QE_489_* = 2000.13, p < 0.001; *marginal R²* = 0.944; *conditional R²* = 0.998). The regression using effective sample-size adjusted standard errors did not detect a significant association between effect size and precision (*slope* = −0.66, *p* = 0.364; *marginal R²* = 0.002; *conditional R²* = 0.932), suggesting that conclusions regarding small-study effects should be interpreted cautiously.

No significant temporal trend in effect sizes was detected (*slope* = −0.037, *p* = 0.133), indicating no evidence that reported effects systematically changed over time. Heterogeneity remained substantial (*QE_489_* = 3229.45, *p* < 0.001; *marginal R²* = 0.026; *conditional R²* = 0.935).

There was evidence of a moderator effect of NMDAR antagonist (*F_(3, 26.05)_* = 56.50, *p* < 0.001). Pooled estimates differed across antagonists, with significant effects observed for ketamine (−2.54, 95% CI [−3.33, −1.75]), MK-801 (−1.56, 95% CI [−2.01, −1.11]), and PCP (−1.88, 95% CI [−2.33, −1.43]) (Figure 3C). Nonetheless, considerable residual heterogeneity persisted after inclusion of NMDAR antagonist as a moderator (*QE_488_* = 3207.037, *p* < 0.001; *marginal R²* = 0.033; *conditional R²* = 0.937). A moderator effect of species was also detected (*F_(3, 24.15)_* = 54.06, *p* < 0.001). Pooled estimates were significant for mice (−2.11, 95% CI [−2.64, −1.57]) and rats (−1.82, 95% CI [−2.16, −1.47]). Although zebrafish were included in the model, the corresponding estimate (−1.77, 95% CI [−3.81, 0.27]) did not reach statistical significance under cluster-robust inference and was based on a small number of studies (Figure 3D). Residual heterogeneity remained present after accounting for species (*QE_488_*= 3215.040, *p* < 0.001; *marginal R²* = 0.005; *conditional R²* = 0.933). A moderator effect of administration schedule was detected (*F_(2, 29.21)_* = 88.99, *p* < 0.001). Significant effects were observed across acute (−1.92, 95% CI [−2.27, −1.57]) and repeated/continuous exposure (−1.89, 95% CI [−2.25, −1.52]) paradigms (Figure S8). Substantial heterogeneity remained after inclusion of administration schedule (*QE_489_* = 3230.051, *p* < 0.001; *marginal R²* = 0.0001; *conditional R²* = 0.932). Effect sizes also differed by developmental stage at induction (*F_(4, 15.86)_* = 39.19, *p* < 0.001). Pooled estimates were significant for adult (−1.98, 95% CI [−2.49, −1.46]), juvenile/adolescent (−1.92, 95% CI [−2.56, −1.27]), and neonatal animals (−1.23, 95% CI [−2.09, −0.37]). Estimates for studies with unclear developmental stage were included in the model but were not interpreted further (Figure S9). Inclusion of the developmental stage as a moderator did not substantially reduce heterogeneity (*QE_487_* = 3011.570, *p* < 0.001; *marginal R²* = 0.008; *conditional R²* = 0.931).

In the multivariate meta-regression including species, NMDAR antagonist, dosing schedule, and developmental stage simultaneously, the overall test of moderators was statistically significant (*F_(9, 26.73)_* = 16.571, *p* < 0.001). The model explained a modest proportion of the between-study heterogeneity (*QE_482_* = 2934.082, *p* < 0.001; *marginal R²* = 0.072; *conditional R²* = 0.942). These findings indicate that, although part of the variability in effect sizes is associated with differences in experimental design characteristics, most heterogeneity remained unexplained after accounting for these moderators, suggesting that additional methodological and/or biological factors likely contribute to variability across studies.

Log-transformed cumulative NMDAR antagonist exposure was available for 453 effect sizes from 171 independent studies with rats and mice and was examined as a continuous moderator (Figure S10). A significant association between cumulative exposure and effect size was detected (*F_(1, 8.29)_* = 8.07, *p* = 0.021). Each ten-fold increase in cumulative exposure was associated with an average change of −0.75 Hedges’ g units (95% CI [−1.36, −0.15] p = 0.021). Considerable unexplained heterogeneity persisted (*QE_451_* = 2780.93, *p* < 0.001; *marginal R²* = 0.165; *conditional R²* = 0.957).

The robustness of the overall pooled effect was examined through a series of sensitivity analyses. Varying the assumed within-experiment correlation resulted in only minor changes in the pooled effect size, with point estimates ranging from −1.95 to −1.85 and overlapping confidence intervals throughout. Leave-one-study-out influence analyses confirmed the stability of the findings; excluding individual studies minimally changed the pooled estimate, all confidence intervals remained statistically significant, and no single study exerted disproportionate influence on the overall result. The overall effect was also re-estimated after excluding studies classified as high overall risk of bias. This analysis, based on 262 effect sizes from 95 independent studies, yielded a pooled effect size of −2.17 (95% CI [−2.61, −1.72]), which was comparable in magnitude to the primary analysis and remained significant. Although heterogeneity remained present (*Q_261_*= 1847.083, *p* < 0.001), the persistence of the effect after the exclusion of high overall risk of bias studies suggests the main findings are robust to study quality considerations.

#### (b) Effects of NMDAR antagonists on locomotor activity measured during social interaction paradigms

All qualitative mappings of study characteristics and the complete results of the analyses conducted for the effects of NMDAR antagonists on locomotor activity measured during social interaction paradigms are available at matheusglls.github.io/NMDA-sb/meta_analysis/nmda_antagonist_effects/locomotion_interaction/n mda_effects_social_interaction_loc.html.

##### (i) Qualitative synthesis

Effect sizes were predominantly derived from rat studies (*k* = 150 from 42 studies), with limited representation from mice (*k* = 17 from 8 studies) and zebrafish (*k* = 3 from 1 study). Among NMDAR antagonists, PCP and MK-801 were most frequently studied (*k* = 78 from 23 studies and *k* = 72 from 24 studies, respectively), whereas ketamine contributed fewer effect sizes (*k* = 20 from 8 studies). The combined distribution across species and antagonists is illustrated in Figure S11A.

##### (ii) Quantitative synthesis

The quantitative synthesis included 170 effect sizes from 51 independent studies. Overall, exposure to NMDAR antagonists was associated with a significant increase in locomotor activity measured during social interaction paradigms when compared with control animals (*Hedges’ g* = 0.40, 95% CI [0.05, 0.75], with a prediction interval of −1.99 to 2.79; Figure S11B). Heterogeneity was high (*I²_total_* = 83.8%), with the majority attributable to between-study variability (*I²_study_* = 71.6%), and a smaller proportion arising from within-study clustering of experiments (*I²_study/exp_* = 12.2%). The test for heterogeneity was significant (*Q_169_* = 992.02, p < 0.001). The estimated variance components were *σ²_study_* = 1.180 and *σ²_study/exp_*= 0.201.

#### (c) Effects of NMDAR antagonists on social preference

All qualitative mappings of study characteristics and the complete results of the analyses conducted for the effects of NMDAR antagonists on social preference are available at matheusglls.github.io/NMDA-sb/meta_analysis/nmda_antagonist_effects/social_preference/nmda_ effects_social_preference.html.

##### (i) Qualitative synthesis

Mice contributed the largest number of effect sizes (*k* = 98 from 56 studies), followed by rats (*k* = 42 from 17 studies), with a smaller number obtained from zebrafish (k = 22 from 8 studies). Across NMDAR antagonists, MK-801 accounted for the largest number of effect sizes (*k* = 71 from 32 studies), followed by ketamine (*k* = 57 from 33 studies) and PCP (*k* = 34 from 18 studies). The distribution of effect sizes across species and NMDAR antagonists is illustrated in Figure 4A, which shows that ketamine and PCP were investigated almost exclusively in rodent models, whereas zebrafish studies were restricted to MK-801 exposure.

**Figure 4.**
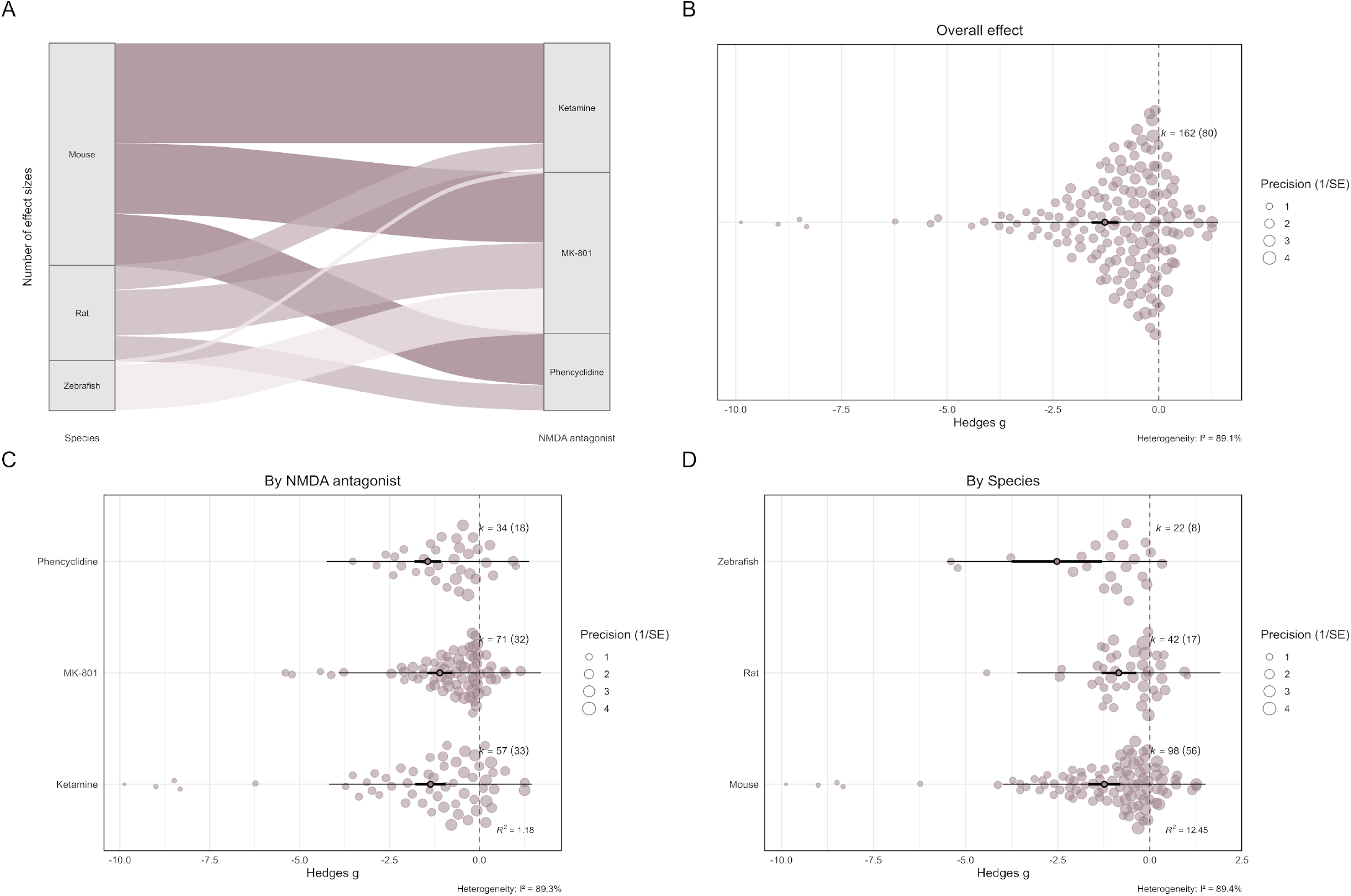
Effects of NMDA receptor antagonists on social preference in laboratory animals. (A) Alluvial plot showing the distribution of effect sizes across species and NMDA antagonists. Flow width represents the number of effect sizes contributing to each category. (B) Orchard plot of the overall pooled effect of NMDA receptor antagonists on social preference. (C) Orchard plots showing effects stratified by NMDA antagonist. (D) Orchard plots showing effects stratified by species. Filled circles represent individual effect sizes scaled by study precision, and open circles indicate pooled point estimates with 95% confidence intervals (thick horizontal lines) and prediction intervals (thin horizontal lines). k denotes the number of effect sizes, with the number of independent studies shown in parentheses.

Intraperitoneal administration was the most frequent route overall (k = 100 from 54 studies; Figures S12–S13) and predominated in mouse and rat studies. Subcutaneous administration was also frequently used (k = 32 from 16 studies), particularly in rodent studies. Other administration routes were sparsely represented, including immersion (k = 22 from 8 studies), which was restricted to zebrafish experiments, minipump delivery (k = 1 from 1 study), and studies in which the route of administration was unclear (k = 7 from 4 studies).

In terms of dosing schedule, acute administration contributed the greatest number of effect sizes (*k* = 88 from 28 studies), whereas repeated dosing was also commonly used (*k* = 71 from 52 studies). Continuous exposure was not frequent (*k* = 3 from 2 studies) and restricted to MK-801 studies (Figure S14).

Studies also varied in the developmental stage at induction. Most effect sizes were derived from adult animals exposed to NMDAR antagonists (*k* = 72 from 31 studies), followed by juvenile/adolescent animals (*k* = 50 from 31 studies). Fewer effect sizes were associated with neonatal exposure (*k* = 12 from 9 studies) or larval stages (*k* = 2 from 1 study). A substantial number of effect sizes originated from studies in which the developmental stage at induction was unclear (*k* = 26 from 9 studies) (Figure S15).

Reporting of sex was uneven across the literature. Most effect sizes were derived from male animals (k = 128, 79.0%; mouse: k = 86; rat: k = 40; zebrafish: k = 2). Female-only samples were rare (k = 4, 2.5%; mouse: k = 4), whereas mixed-sex samples contributed a smaller proportion of the evidence (k = 23, 14.2%; mouse: k = 5; zebrafish: k = 18). A limited number of effect sizes were derived from studies in which sex was unclear (k = 7, 4.3%; mouse: k = 3; rat: k = 2; zebrafish: k = 2) (Figure S16).

##### (ii) Quantitative synthesis

The quantitative synthesis comprised 162 effect sizes derived from 80 independent studies. Overall, exposure to NMDAR antagonists was associated with a reduction in social preference compared with animals in the control group (*Hedges’ g* = −1.27, 95% CI [−1.59, −0.95], with a prediction interval of −3.95 to 1.41; Figure 4B). Heterogeneity was high (*I²_total_*= 89.1%). The majority of this heterogeneity was attributable to between-study variability (*I²_study_* = 88.2%), with only a minimal contribution from within-study clustering (*I²_study/exp_* = 1.0%). The test for heterogeneity was significant (*Q_161_* = 719.16, p < 0.001), confirming that observed variation in effect sizes exceeded what would be expected from sampling error alone. Variance components further supported this pattern, with substantial variance at the study level (*σ²_study_* = 1.77) and comparatively little variance attributable to clustering of multiple effect sizes within experiments (*σ²_study/exp_*= 0.02).

Visual inspection of the funnel plots suggested marked asymmetry, consistent with small-study effects, which may reflect publication bias (Figure S17). PET indicated a significant association between effect size and study precision (*slope* = −5.63, *p* < 0.001; *marginal R²* = 0.802; *conditional R²* = 0.938), providing strong evidence of small-study effects. The PEESE model also identified a significant association between effect size and sampling variance (*slope* = −3.50, *p* < 0.001). The bias-adjusted intercept from the PEESE was small and not statistically significant (Hedges’ g = 0.04, 95% CI [−0.21, 0.29]), indicating that adjustment for small-study effects eliminated the statistical significance of the overall effect estimate. Residual heterogeneity remained high following bias adjustment (*QE_160_* = 390.59, *p* < 0.001; *marginal R²* = 0.866; *conditional R²* = 0.984). Using effective-sample-size adjusted standard errors, the regression did not identify a significant association between effect size and precision (*slope* = −0.35, *p* = 0.625; *marginal R²* = 0.001; *conditional R²* = 0.982), suggesting that evidence for small-study effects should be interpreted with caution.

No evidence of a temporal trend in effect sizes was observed. Meta-regression of effect size on publication year revealed no significant association (*slope* = 0.01, p = 0.785). Substantial residual heterogeneity persisted in the time-lag model (*QE_160_* = 712.51, p < 0.001; *marginal R²* = 0.001; *conditional R²* = 0.989).

There was evidence of a moderator effect of NMDAR antagonist (*F_(3, 4.14)_* = 18.83, *p* = 0.007). Pooled estimates differed across antagonists, with significant reductions in social preference observed for ketamine (*Hedges’ g* = −1.36, 95% CI [−1.79, −0.94]), MK-801 (−1.10, 95% CI [−1.45, −0.75]), and PCP (−1.44, 95% CI [−1.87, −1.01]) (Figure 4C). However, substantial residual heterogeneity remained after accounting for NMDAR antagonist (*QE_159_* = 713.960, *p* < 0.001; *marginal R²* = 0.012; *conditional R²* = 0.984). A moderator effect of species was also detected (*F_(3, 14.29)_* = 19.81, *p* < 0.001) (Figure 4D). Pooled estimates were significant for mice (−1.24, 95% CI [−1.67, −0.80]), rats (−0.84, 95% CI [−1.34, −0.34]), and zebrafish (−2.52, 95% CI [−4.16, −0.89]); however, the latter was based on a relatively small number of studies and has wide confidence intervals, and should therefore be interpreted with caution. Inclusion of species as a moderator reduced heterogeneity attributable to within-study clustering, but residual heterogeneity remained significant (*QE_159_* = 696.651, *p* < 0.001; *marginal R²* = 0.124; *conditional R²* = 1.000), suggesting that species explained only a limited proportion of the overall variability. A moderator effect of administration schedule was detected (*F_(2, 14.93)_* = 32.52, *p* < 0.001). Significant effects were observed across acute (−1.00, 95% CI [−1.51, −0.49]) and repeated/continuous exposure paradigms (−1.41, 95% CI [−1.84, −0.98]) (Figure S18). Substantial heterogeneity remained after including the administration schedule as a moderator (*QE*_160_ = 610.659, *p* < 0.001; *marginal R²* = 0.026; *conditional R²* = 0.962). Effect sizes also differed by developmental stage at induction (*F_(4, 16.02)_* = 14.87, *p* < 0.001). Pooled estimates were significant for adult animals (−1.35, 95% CI [−1.80, −0.89]) and juvenile/adolescent animals (−1.33, 95% CI [−1.97, −0.68]). The estimate for neonatal/larval animals was also negative but did not reach statistical significance under cluster-robust inference (−1.22, 95% CI [−2.48, 0.04]). Estimates for studies with unclear developmental stage were included in the model but were not interpreted further (Figure S19). Inclusion of the developmental stage as a moderator did not substantially reduce heterogeneity (*QE_158_* = 702.942, *p* < 0.001; *marginal R²* = 0.009; *conditional R²* = 0.990).

When species, NMDAR antagonist, dosing schedule, and developmental stage were included simultaneously in a multivariate meta-regression model, the overall set of moderators was statistically significant (*F_(9, 5.39)_* = 4.99, *p* = 0.040). Despite this, the proportion of heterogeneity accounted for by these variables was limited (*QE_153_* = 555.237, *p* < 0.001; *marginal R²* = 0.158; *conditional R²* = 0.993), indicating that most between-study variability remained unexplained.

Log-transformed cumulative NMDAR antagonist exposure was available for 135 effect sizes from 68 independent studies conducted in rats and mice (Figure S20). No significant association between cumulative exposure and effect size was detected (*F_(1, 3.83)_* = 7.24, p = 0.057). Although the estimated slope was negative, indicating larger reductions in social preference with increasing exposure (−0.24 Hedges’ g units per ten-fold increase; 95% CI [−0.50, 0.01]), this association did not reach statistical significance under cluster-robust inference. Substantial unexplained heterogeneity remained after inclusion of cumulative exposure as a moderator (*QE_133_* = 505.06, *p* < 0.001; *marginal R²* = 0.057; *conditional R²* = 0.989).

Varying the assumed within-experiment correlation resulted in only minimal changes in the pooled effect size, with point estimates ranging from −1.28 to −1.26 and highly overlapping confidence intervals throughout. Leave-one-study-out influence analyses confirmed the stability of the findings; excluding individual studies minimally changed the pooled estimate, and all confidence intervals remained statistically significant. When studies judged to be at high overall risk of bias were excluded, the meta-analysis comprised 82 effect sizes from 50 independent studies and yielded a pooled effect of −1.22 (95% CI [−1.59, −0.85]). This estimate was similar in magnitude to the primary analysis and remained significant. Despite this, heterogeneity remained present (Q81 = 371.19, p < 0.001).

#### (d) Effects of NMDAR antagonists on locomotor activity measured during social preference paradigms

All qualitative mappings of study characteristics and the complete results of the analyses conducted for the effects of NMDAR antagonists on locomotor activity measured during social preference paradigms are available at matheusglls.github.io/NMDA-sb/meta_analysis/nmda_antagonist_effects/locomotion_preference/nmda_effects_social_preference_loc.html.

##### (i) Qualitative synthesis

Most effect sizes were derived from mouse studies (*k* = 24 from 6 independent studies), with relatively few from rats (*k* = 5 from 2 studies) and zebrafish (*k* = 5 from 3 studies). Across NMDAR antagonists, MK-801 was most frequently examined (*k* = 18 from 8 studies), whereas PCP (*k* = 9 from 2 studies) and ketamine (*k* = 7 from 3 studies) were less commonly represented. These distributions are summarised in Figure S21A.

#### (ii) Quantitative synthesis

The quantitative synthesis included 34 effect sizes derived from 11 independent studies. Overall, exposure to NMDAR antagonists was not associated with a statistically significant change in locomotor activity measured during the social preference test (*Hedges’ g* = 0.18, 95% CI [−0.17, 0.53], with a prediction interval ranging from −0.62 to 0.99; Figure S21B). Heterogeneity was moderate (*I²_total_* = 34.4%), with most of the variability attributable to between-study differences (*I²_study_* = 26.8%) and a smaller proportion arising from within-study clustering of experiments (*I²_study/exp_* = 7.6%). The test for heterogeneity was statistically significant (*Q_33_* = 132.27, *p* < 0.001). The estimated variance components were *σ²_study_*= 0.076 and *σ²_study/exp_* = 0.021.

#### (e) Effects of antipsychotics on NMDAR antagonist–induced alterations in social interaction

All qualitative mappings of study characteristics and the complete results of the analyses conducted for the effects of antipsychotics on NMDAR antagonist–induced alterations in social interaction are available at matheusglls.github.io/NMDA-sb/meta_analysis/antipsychotics_on_nmda/social_interaction/atp_effe cts_social_interaction.html.

##### (i) Qualitative synthesis

Rats contributed the majority of effect sizes (*k* = 190 from 42 independent studies), followed by mice (*k* = 58 from 24 studies). PCP accounted for the largest share of NMDAR antagonist–related effect sizes (*k* = 140 from 33 studies), followed by MK-801 (*k* = 72 from 27 studies), while ketamine contributed comparatively fewer effect sizes (*k* = 36 from 14 studies). A wide range of antipsychotics was investigated, with clozapine contributing the largest number of effect sizes (*k* = 70 from 33 studies), followed by haloperidol (*k* = 50 from 21 studies), risperidone (*k* = 39 from 17 studies), and olanzapine (*k* = 28 from 13 studies). Other antipsychotics, including aripiprazole (*k* = 20 from 8 studies), quetiapine (*k* = 8 from 1 study), sulpiride (*k* = 8 from 3 studies), ziprasidone (*k* = 6 from 3 studies), amisulpride (*k* = 5 from 2 studies), lurasidone (*k* = 5 from 2 studies), chlorpromazine (*k* = 3 from 2 studies), and cariprazine (*k* = 6 from 2 studies), were represented by a smaller number of effect sizes. The distribution of effect sizes across species, NMDAR antagonists, and antipsychotic drugs is illustrated in Figure 5A.

**Figure 5.**
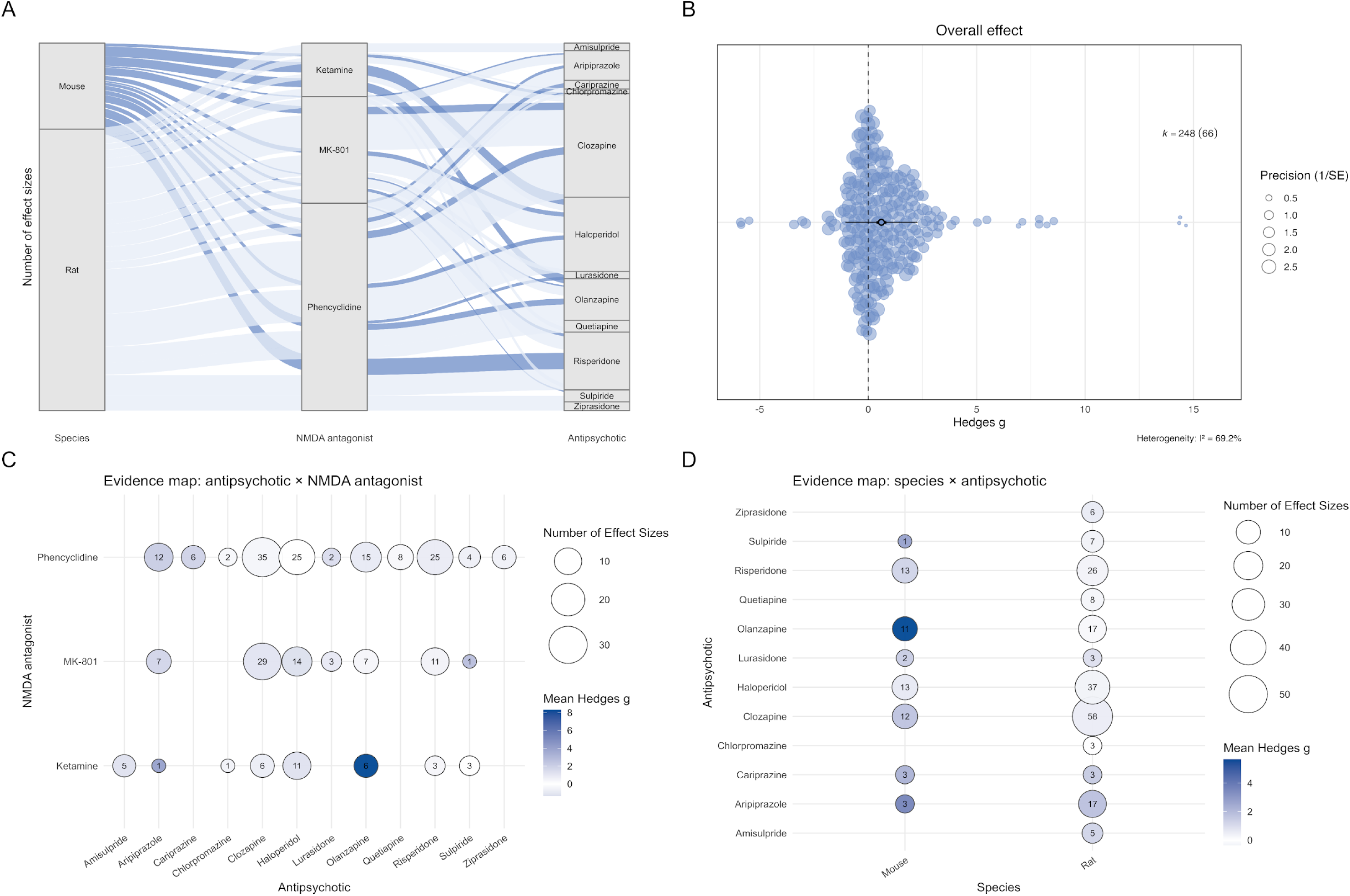
Effects of antipsychotics on NMDA receptor antagonist–induced alterations in social interaction in laboratory animals. (A) Alluvial plot showing the distribution of effect sizes across species, NMDA receptor antagonists, and antipsychotics. Flow width represents the number of effect sizes contributing to each category. (B) Orchard plot of the overall pooled effect of antipsychotics on NMDA antagonist–induced changes in social interaction. Filled circles represent individual effect sizes scaled by study precision, and the open circle indicates the pooled point estimate with 95% confidence intervals (thick horizontal lines) and prediction interval (thin horizontal lines). k denotes the number of effect sizes, with the number of independent studies shown in parentheses. (C) Evidence map of effect sizes by NMDA antagonist and antipsychotic. Circle size represents the number of effect sizes, and colour intensity indicates the mean Hedges’ g. (D) Evidence map of effect sizes by species and antipsychotic, with circle size indicating the number of effect sizes and colour representing the mean Hedges’ g.

For PCP, the most frequently represented pairings with antipsychotics were with clozapine (*k* = 35), haloperidol (*k* = 25), and risperidone (*k* = 25) (Figure 5C). For MK-801, most estimates involved clozapine (*k* = 29) and haloperidol (*k* = 14). Ketamine-related estimates were less common, with the largest numbers reported for haloperidol (k = 11), followed by clozapine and olanzapine (each k = 6); most other ketamine–antipsychotic pairings were represented by only a small number of effect sizes.

Across antipsychotics, most effect sizes were derived from studies using rats, with fewer estimates available in mice (Figure 5D). In rats, clozapine (*k* = 58), haloperidol (*k* = 37), risperidone (*k* = 26), and olanzapine (*k* = 17) contributed the largest numbers of effect sizes. In mice, the highest representation was for haloperidol (*k* = 13), risperidone (*k* = 13), and olanzapine (*k* = 11), with fewer effect sizes available for the remaining compounds.

##### (ii) Quantitative synthesis

The quantitative synthesis included 248 effect sizes derived from 66 independent studies. Overall, antipsychotic treatment was associated with increased social interaction in NMDAR antagonist–exposed animals (*Hedges’ g* = 0.61, 95% CI [0.38, 0.83], with a prediction interval ranging from −1.06 to 2.27; Figure 5B). Heterogeneity was substantial (*I²_total_* = 69.2%), with nearly half of the variability attributable to between-study differences (*I²_study_*= 48.0%) and a further proportion arising from within-study clustering of experiments (*I²_study/exp_* = 21.2%). The test for heterogeneity was statistically significant (*Q_247_* = 1191.89, *p* < 0.001). The estimated variance components were *σ²_study_* = 0.471 and *σ²_study/exp_* = 0.208.

#### (f) Effects of antipsychotics on locomotor activity measured during social interaction paradigms in animals exposed to NMDAR antagonists

All qualitative mappings of study characteristics and the complete results of the analyses conducted for the effects of antipsychotics on locomotor activity measured during social interaction paradigms in animals exposed to NMDAR antagonists are available at matheusglls.github.io/NMDA-sb/meta_analysis/antipsychotics_on_nmda/locomotion_interaction/at p_effects_social_interaction_loc.html.

##### (i) Qualitative synthesis

Studies using rats contributed almost all effect sizes (k = 121 from 18 independent studies), with only one effect size obtained from mice (k = 1 from 1 study). PCP accounted for the largest number of effect sizes (k = 69 from 11 studies), followed by MK-801 (k = 44 from 7 studies), while ketamine contributed comparatively fewer effect sizes (k = 9 from 4 studies). A range of antipsychotics was examined, with clozapine contributing the largest number of effect sizes (k = 39 from 11 studies), followed by haloperidol (k = 29 from 7 studies), risperidone (k = 21 from 7 studies), and olanzapine (k = 9 from 3 studies). Other antipsychotics, including aripiprazole (k = 7 from 2 studies), quetiapine (k = 8 from 1 study), sulpiride (k = 4 from 1 study), chlorpromazine (k = 2 from 1 study), and cariprazine (k = 3 from 1 study), were represented by a smaller number of effect sizes. The distribution of effect sizes across species, NMDAR antagonists, and antipsychotic drugs is illustrated in Figure S22A.

##### (ii) Quantitative synthesis

The quantitative synthesis included 122 effect sizes derived from 19 independent studies. Overall, antipsychotic treatment was associated with a significant reduction in locomotor activity measured during social interaction paradigms when compared to animals treated with NMDAR antagonists alone (*Hedges’ g* = −0.49, 95% CI [−0.70, −0.29], with a prediction interval ranging from −1.31 to 0.32; Figure S22B). Heterogeneity was moderate (*I²_total_* = 33.7%), with variability arising from both between-study differences (*I²_study_*= 11.0%) and within-study clustering of experiments (*I²_study/exp_*= 22.7%). The test for heterogeneity was statistically significant (*Q_121_*= 678.70, *p* < 0.001). The estimated variance components were *σ²_study_*= 0.044 and *σ²_study/exp_* = 0.090.

#### (g) Effects of antipsychotics on NMDAR antagonist–induced alterations in social preference

All qualitative mappings of study characteristics and the complete results of the analyses conducted for the effects of antipsychotics on NMDAR antagonist–induced alterations in social preference are available at matheusglls.github.io/NMDA-sb/meta_analysis/antipsychotics_on_nmda/social_preference/atp_eff ects_social_preference.html.

##### (i) Qualitative synthesis

Mice contributed the majority of effect sizes (*k* = 29 from 17 independent studies), followed by rats (*k* = 8 from 3 studies), with a small number obtained from zebrafish (*k* = 3 from 1 study). Ketamine accounted for the largest number of effect sizes (*k* = 20 from 10 studies), followed by MK-801 (*k* = 14 from 6 studies), while PCP contributed fewer effect sizes overall (*k* = 6 from 5 studies). Several antipsychotics were investigated, with risperidone contributing the largest number of effect sizes (*k* = 14 from 10 studies), followed by clozapine (*k* = 7 from 6 studies), aripiprazole (*k* = 6 from 4 studies), olanzapine (*k* = 6 from 5 studies), and haloperidol (*k* = 5 from 4 studies). Other antipsychotics, including sulpiride (*k* = 1 from 1 study) and amisulpride (*k* = 1 from 1 study), were represented by a single effect size each. The distribution of effect sizes across species, NMDAR antagonists, and antipsychotic drugs is illustrated in Figure 6A.

**Figure 6.**
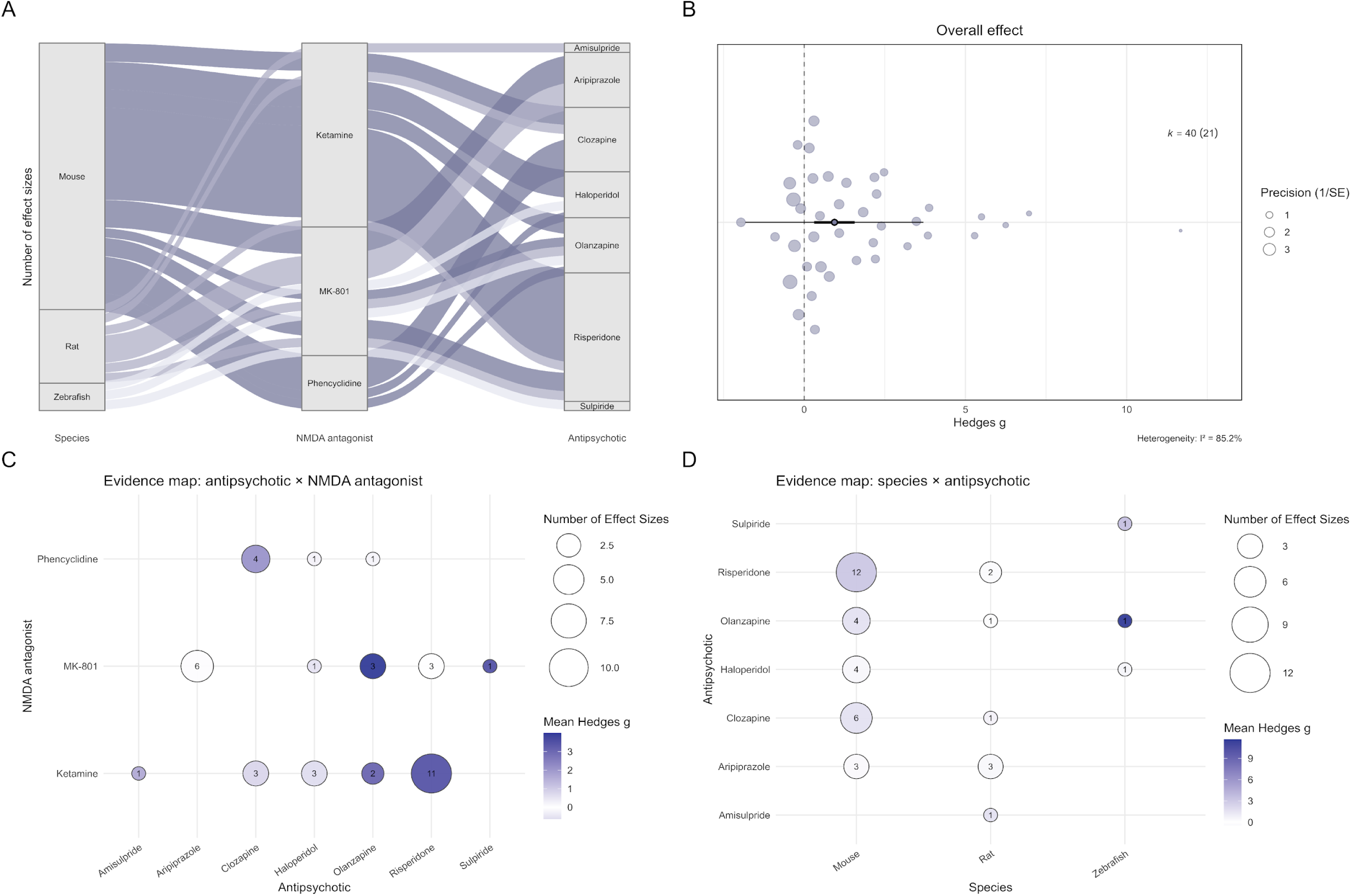
Effects of antipsychotics on NMDA receptor antagonist–induced alterations in social preference in laboratory animals. (A) Alluvial plot showing the distribution of effect sizes across species, NMDA receptor antagonists, and antipsychotics. Flow width represents the number of effect sizes contributing to each category. (B) Orchard plot of the overall pooled effect of antipsychotics on NMDA antagonist–induced changes in social preference. Filled circles represent individual effect sizes scaled by study precision, and the open circle indicates the pooled point estimate with 95% confidence intervals (thick horizontal lines) and prediction interval (thin horizontal lines). k denotes the number of effect sizes, with the number of independent studies shown in parentheses. (C) Evidence map of effect sizes by NMDA antagonist and antipsychotic. Circle size represents the number of effect sizes, and colour intensity indicates the mean Hedges’ g. (D) Evidence map of effect sizes by species and antipsychotic, with circle size indicating the number of effect sizes and colour representing the mean Hedges’ g.

Within the ketamine studies, risperidone was the most frequently reported antipsychotic (*k* = 11), with fewer effect sizes for clozapine (*k* = 3), haloperidol (*k* = 3), olanzapine (*k* = 2), and a single estimate for amisulpride (*k* = 1) (Figure 6C). Under MK-801, most estimates involved aripiprazole (*k* = 6), followed by olanzapine (*k* = 3) and risperidone (*k* = 3), with single estimates for haloperidol (*k* = 1) and sulpiride (*k* = 1). PCP contributed comparatively few estimates overall and was largely represented by clozapine (*k* = 4), with single estimates for haloperidol (*k* = 1) and olanzapine (*k* = 1).

Across species, mouse studies contributed the bulk of the evidence for social preference outcomes, particularly for risperidone (*k* = 12), alongside clozapine (*k* = 6), haloperidol (*k* = 4), olanzapine (*k* = 4), and aripiprazole (*k* = 3) (Figure 6D). Rat studies contributed fewer effect sizes, mainly for aripiprazole (*k* = 3) and risperidone (*k* = 2), with single estimates for clozapine (*k* = 1), olanzapine (*k* = 1), and amisulpride (*k* = 1). Zebrafish data were limited to single estimates for olanzapine, haloperidol, and sulpiride (*k* = 1 each).

##### (ii) Quantitative synthesis

The quantitative synthesis included 40 effect sizes derived from 21 independent studies. Overall, antipsychotic treatment was associated with a significant increase in social preference in animals exposed to NMDAR antagonists (*Hedges’ g* = 0.94, 95% CI [0.31, 1.57], with a prediction interval ranging from −1.82 to 3.70; Figure 6B). Heterogeneity was high (*I²_total_*= 85.2%), with all observed variability attributable to between-study differences (*I²_study_* = 85.2%), and no additional contribution from within-study clustering of experiments (*I²_study/exp_* = 0%). The test for heterogeneity was statistically significant (*Q_39_*= 353.19, *p* < 0.001). The estimated variance components were *σ²_study_* = 1.66 and *σ²_study/exp_* = 0.

## IV. DISCUSSION

Across the included studies, exposure to NMDAR antagonists was associated with reduced social behaviour overall; however, the robustness of this pattern varied depending on the behavioural paradigm and on bias diagnostics. In social interaction tests, NMDAR antagonists significantly reduced social interaction relative to controls, with substantial heterogeneity and wide prediction intervals; although the effect size was attenuated after adjustment for small-study effects, it remained statistically significant. For social preference paradigms, the pooled estimate was also significant, yet this effect was eliminated after bias adjustment, indicating greater sensitivity to imprecise studies and potential publication bias. Locomotor activity assessed during social paradigms provided important interpretive context, as antagonists increased activity during social interaction tests but showed no clear effects during social preference. Finally, clinically approved antipsychotics were associated with higher social interaction and social preference in animals exposed to NMDAR antagonists, with reduced locomotion in social interaction paradigms, indicating that apparent normalisation of social outcomes co-occurred with changes in general activity.

In this systematic review, we analysed social interaction and social preference as distinct outcome domains, not only because they operationalise social behaviour in different ways, but also because the evidence bases underpinning each domain have different profiles. Social interaction outcomes were drawn from a much larger evidence base and covered a longer publication period (1989 to 2023, median 2013), whereas social preference outcomes came from a smaller and more recent literature (2009 to 2023, median 2016). Although these differences in publication era and reporting practices could, in principle, contribute to discrepancies between domains, the time-lag analyses showed no systematic association between effect size and publication year in either domain, suggesting that simple temporal trends are unlikely to explain the contrast between paradigms.

Both domains were represented across the same species, including rats, mice, and zebrafish, but the distribution of studies across species and interventions differed between paradigms. In the extracted dataset, social interaction outcomes predominantly comprised studies with rats and administration of PCP or MK-801, and most frequently employed repeated dosing designs alongside acute exposures to antagonists. Social preference outcomes relied more heavily on mice, included a proportionally larger contribution from zebrafish studies, and were informed by a relatively greater share of acute paradigms. Even with this methodological diversity, our results are consistent with the view that NMDAR antagonism reduces social behaviour, whether measured as reciprocal interaction or as social preference (Neill *et al*., 2010, 2014; Lee & Zhou, 2019). However, although the average effects in both domains pointed towards reduced sociability, the prediction intervals indicate that future studies could plausibly observe anything from reduced sociability to no detectable change and, in some contexts, increased sociability. Consistent with this, the literature includes both social interaction and social preference studies reporting little or no change (Becker & Grecksch, 2004; Egerton *et al*., 2008; Seillier & Giuffrida, 2016; Herzog *et al*., 2020) as well as studies reporting increased sociability following NMDAR antagonist exposure (Dunn *et al*., 1989; Jenkins *et al*., 2008; Rawat *et al*., 2022; Giongo *et al*., 2023).

This combination of methodological diversity and wide prediction intervals requires careful interpretation, with particular emphasis on bias diagnostics (Riley, Higgins & Deeks, 2011; Sterne *et al*., 2011; Stanley & Doucouliagos, 2014; Zwetsloot *et al*., 2017). Concerns about inflated effect estimates and the selective publication of clear behavioural findings are well documented in preclinical animal research (Sena *et al*., 2010; Korevaar, Hooft & Ter Riet, 2011; Macleod *et al*., 2015). In our data, this issue was particularly evident for social preference. Although the conventional pooled estimate suggested reduced preference, this signal was not robust to adjustment for small-study effects, and the estimate was compatible with no effect. By contrast, the social interaction effect remained significant after adjustment for small-study effects, suggesting a more robust signal in that domain. Notably, regressions using effective-sample-size adjusted standard errors were not statistically significant for either primary outcome, indicating that evidence for small-study effects was not fully consistent across alternative modelling approaches and should therefore be interpreted cautiously. However, small-study effects are only one pathway through which effect sizes can be inflated in preclinical research (Button *et al*., 2013; Munafò *et al*., 2017). Low statistical power and selective reporting can exaggerate observed effects, with analytic flexibility yielding overconfident inferences even when individual decisions appear defensible (Simmons, Nelson & Simonsohn, 2011; Gelman & Loken, 2013; Button *et al*., 2013). This concern is relevant to the present dataset, as extracted experiments generally used small sample sizes, typically below 10 animals per experimental group. In addition, incomplete reporting of key design and analytical features can make it difficult to distinguish bias from genuine heterogeneity across laboratories and protocol families (Landis *et al*., 2012; Percie du Sert *et al*., 2020). Taken together, these issues lead to cautious interpretation of the magnitude and generalisability of pooled effects, including those that remain significant after bias adjustment.

These concerns are amplified by the way social behaviour is operationalised in the primary literature. In the included studies, social interaction was most commonly assessed by measuring total interaction time, often complemented by measures of direct contact or investigatory behaviours. For social preference, sociability was most commonly indexed by time spent in a social zone or compartment. However, the operational definition of interaction and related endpoints varies significantly across studies, resulting in ambiguity regarding the observed variables and limiting comparability across experiments. This diversity raises the possibility that different studies are quantifying distinct behavioural components that are not directly comparable and may not carry the same ecological relevance or pharmacological sensitivity, even when they are discussed as evidence of a single social deficit (Saverino C & Gerlai R, 2008; Millan *et al*., 2014; Arakawa, 2018; Kondrakiewicz *et al*., 2019; Geng & Peterson, 2019; Ogi A *et al*., 2020). A more comprehensive mapping of relevant social outcomes remains needed, alongside explicit specification of which component of social behaviour is being targeted in a given paradigm or outcome. Apart from that, consistent reporting of primary outcomes would help limit selective emphasis on favourable endpoints and reduce the scope for *post hoc* rationalisation, when multiple measures are available, a concern that has also been documented in behavioural neuroscience (Simmons *et al*., 2011; Rosso *et al*., 2024).

Moderator analyses were used to test whether variation in effects across studies was consistent with identifiable experimental and biological differences and to provide a partial explanation for the substantial heterogeneity observed in both domains (Yang *et al*., 2023). The moderator results were broadly in line with the literature by showing reduced social behaviour across NMDAR antagonists, species, schedule, and developmental stages (Neill *et al*., 2010, 2014; Millan *et al*., 2014; Lee & Zhou, 2019). For zebrafish, the interaction estimate was imprecise and did not remain significant under robust inference. Given the small number of zebrafish studies and the diversity of paradigms and designs used in this subset, this pattern is more consistent with limited evidence and heterogeneity than with a reliable species difference (Benvenutti *et al*., 2022).

Even after accounting for these moderators, both in analyses evaluating individual moderators separately and in multivariable models combining multiple experimental characteristics simultaneously, heterogeneity remained high in both social interaction and social preference, suggesting that important sources of between-study variation were not captured by the moderators that could be analysed with adequate support. This is not unexpected in preclinical evidence synthesis, where studies are often small and highly diverse, and differences in species, dosing regimens, testing context, and endpoint definitions can generate wide dispersion in effect sizes (Hooijmans *et al*., 2014a; Vesterinen *et al*., 2014; Yang *et al*., 2023; Ineichen *et al*., 2024b). In our dataset, incomplete reporting and sparse data within many candidate moderator levels limited the ability to examine additional sources of heterogeneity with confidence, consistent with broader evidence that deficiencies in reporting are common in animal research and constrain what systematic reviews can explain (Leung *et al*., 2018; Hunniford *et al*., 2021; Ritskes-Hoitinga & Pound, 2022; Wilson *et al*., 2023).

To examine whether dosing intensity contributed to variation in effect sizes and residual heterogeneity, we modelled cumulative exposure in rodents, as dosing information was sufficiently consistent in rat and mouse studies. Higher cumulative exposure to NMDAR antagonists was associated with larger reductions in social interaction, while the corresponding analysis in social preference did not show a clear association in our dataset. This direction is consistent with reviews of rodent NMDAR hypofunction models, which describe repeated or subchronic antagonist regimens as a way to produce more sustained behavioural phenotypes, including social withdrawal (Neill *et al*., 2014; Lee & Zhou, 2019). Direct within-study comparisons of acute versus repeated or chronic exposure, or of different exposure durations, are rare in the social behaviour literature, and regimen-related differences are therefore often inferred across studies. Across the small subset of studies that directly compare exposure regimens within the same experiment, findings remain mixed, with some studies suggesting regimen or timing sensitivity and others reporting little or no difference between acute and repeated schedules (Sams-Dodd, 1996; Becker & Grecksch, 2004; Li, He & Munro, 2012; Hanks *et al*., 2013; McKibben, Reynolds & Jenkins, 2014; Cieślik *et al*., 2019). Accordingly, the evidence base still lacks adequately powered head-to-head comparisons that systematically manipulate regimen parameters across NMDAR antagonists, leaving an important gap in the social behaviour literature.

Hyperlocomotion is a well-documented feature of NMDAR antagonist exposure in preclinical models and is commonly interpreted as a phenotype with relevance to the positive symptoms of schizophrenia (Large, 2007; Neill *et al*., 2010, 2014; Frohlich & Van Horn, 2014; Lee & Zhou, 2019; Benvenutti *et al*., 2022). In the social interaction studies, NMDAR antagonists were associated with higher locomotor activity measured during the test, while reciprocal social engagement was reduced. This is relevant for interpreting social interaction assays because the test sums active social behaviour over a brief encounter in a novel arena; thus, scores depend on how animals allocate time between social contact, exploration, and other competing activities during the session (File & Hyde, 1978; File & Seth, 2003; Toth & Neumann, 2013; Kondrakiewicz *et al*., 2019). More broadly, the potential for non-social behavioural changes to shape apparent social effects is not unique to NMDAR antagonists and has been explored in the context of psychostimulant (Deak *et al*., 2009; Šlamberová *et al*., 2010, 2015) and sedative (File & Seth, 2003) drugs and across behavioural models where altered arousal, exploration, or motor output can shift time budget-based social endpoints (Nadler *et al*., 2004; Silverman *et al*., 2010; Yang *et al*., 2011). In NMDAR antagonist models, reduced social interaction alongside increased activity makes a simple global behavioural suppression account unlikely, being consistent with drug-induced changes in how behaviour is organised during the encounter, including more competing non-social exploration and shorter or more fragmented affiliative bouts. Accordingly, stronger inference would come from routinely pairing social outcomes with within-task locomotion and complementary ethological measures such as bout structure and partner-directed investigation and from explicitly distinguishing acute drug state effects from delayed or post-drug phenotypes when claims about social dysfunction or pharmacological rescue are made (File & Hyde, 1978; File & Seth, 2003; Nadler *et al*., 2004; Silverman *et al*., 2010; Yang *et al*., 2011; Peleh *et al*., 2019; Jabarin, Netser & Wagner, 2022).

In the social preference literature, locomotor activity did not show a clear overall effect in our synthesis, but this does not imply that non-social influences are insignificant. Preference readouts are fundamentally time budget measures of where an animal spends time in the apparatus (Nadler *et al*., 2004; Yang *et al*., 2011). Small shifts in thigmotaxis, side bias, or avoidance of exposed zones can alter chamber or zone occupancy and therefore alter derived sociability indices even when overall distance travelled is not consistently different across groups (Yang *et al*., 2011; Kondrakiewicz *et al*., 2019). For this reason, social preference protocols recommend reporting activity proxies such as chamber entries and exploration measures alongside occupancy-based sociability indices (Nadler *et al*., 2004; Yang *et al*., 2011), yet these measures are not consistently collected or reported; focusing on one or two preference metrics can miss non-social drivers of performance that affect interpretation (Kondrakiewicz *et al*., 2019; Winiarski *et al*., 2022; Jabarin *et al*., 2022). Taken together, the absence of a clear locomotor signal in this subset may reflect both genuine task-specific expression of the phenotype and the smaller, potentially selective evidence base for within-task activity reporting in social preference studies.

Antipsychotic treatment was associated with increased social behaviour, but the accompanying locomotor pattern suggests that this should not be interpreted as selective restoration of social function in isolation. Because social interaction measures depend on animals being active in the arena (File & Hyde, 1978; File & Seth, 2003), the co-occurrence of higher social behaviour with reduced locomotor activity suggests that at least part of the apparent rescue may reflect a change in time allocation and reduced behavioural competition during the encounter rather than a selective change in social processes. This interpretation is also consistent with the broader point that many antipsychotics can produce sedation and reduced activity in humans (Hynes *et al*., 2020; Nomura *et al*., 2025) and laboratory animals (Bernardi, De Souza & Palermo Neto, 1981; van der Zwaal *et al*., 2010; McOmish *et al*., 2012; Lian *et al*., 2015), so state effects are a plausible contributor to time-based social outcomes when activity shifts in parallel. The antipsychotic literature is also concentrated in a small number of compounds, most commonly clozapine, haloperidol, risperidone, and olanzapine.

These findings should be interpreted alongside a previous systematic review and meta-analysis of treatments for social interaction impairments in animal models of schizophrenia, which also reported beneficial treatment effects, particularly for atypical antipsychotics (Hazani, Lavidor & Weller, 2022). However, important differences in scope and design limit direct comparison. That review synthesised treatment effects across multiple schizophrenia-related models and treatment classes, included only studies in which the model produced a social interaction deficit, and restricted outcomes to social interaction time or number of interactions. This design is appropriate for asking whether treatments reverse an established deficit, but it may favour evidence in which the expected phenotype and treatment response are already present, thereby overemphasising pharmacological responsiveness as a marker of model validity. It also does not distinguish between social interaction and social preference paradigms or evaluate locomotor activity, both of which are central to interpreting social outcomes. Our results are consistent with antipsychotic-related increases in social behaviour, but extend previous evidence by placing these effects within a more specific NMDAR antagonist framework and by showing that apparent social improvement should be interpreted cautiously.

None of this excludes pharmacologically meaningful effects on social behaviour, because antipsychotics may increase the likelihood that animals approach and maintain contact with social stimuli. Even if such effects are genuine in the assay, the modest and inconsistent clinical signal for primary negative symptoms and social dysfunction means that apparent preclinical rescue should still not be overinterpreted (Correll & Schooler, 2020; Spark *et al*., 2022). Ultimately, basic measures of social proximity in laboratory animals may lack construct validity to meaningfully reflect the complex negative symptoms experienced by patients with schizophrenia. This caution fits with wider meta-research on translation, which highlights gaps between preclinical and clinical practice, concerns about preclinical study rigour, and attenuation from animal to human effects, with only a small fraction of animal-tested interventions reaching regulatory approval (Ineichen *et al*., 2024a). It also aligns with translational critiques from other disease areas, where even apparently clinically relevant outcomes can have limited translational validity and where a narrow set of experimental practices can dominate the evidence base (Berg *et al*., 2024). In this context, the antipsychotic evidence mainly shows that the assays are pharmacologically sensitive, while the link between changes in preclinical social endpoints and clinically meaningful improvements in social functioning remains uncertain. More broadly, our findings challenge the view that NMDAR antagonist–based social behaviour assays provide strong predictive validity for the negative symptom domain of schizophrenia. Although these assays detect social behavioural changes and are responsive to clinically approved antipsychotics, this pattern does not establish that they capture clinically meaningful social dysfunction. Instead, the apparent translational gap may reflect not only true cross-species or mechanistic failure, but also the cumulative effects of publication bias, heterogeneity, limited construct validity, and non-specific behavioural changes within the preclinical evidence base.

Risk of bias assessment highlighted a familiar pattern in *in vivo* neuroscience, where key safeguards are often ignored or reported too vaguely to support confident judgment, particularly for randomisation, allocation concealment, and blinded outcome assessment (Worp *et al*., 2010; Macleod *et al*., 2015). Although excluding studies rated at high overall risk did not materially change the main findings, this sensitivity analysis has clear limits when most judgements fall into an “unclear” category rather than “low” risk. In that situation, the key issue is not simply a small subset of demonstrably flawed studies, but the possibility that important design and analysis practices are inconsistently implemented and selectively reported. The high prevalence of “unclear” ratings, therefore, primarily reflects reporting limitations and reinforces the need for better transparency in preclinical methods and analysis. This is particularly relevant because efforts to improve reporting quality, including guideline-based initiatives, have not consistently delivered large improvements in practice (Leung *et al*., 2018; Hair *et al*., 2019), so risk of bias assessment will remain constrained unless reporting standards are more reliably adopted and enforced.

This methodological uncertainty is also evident in the inter-rater reliability results; agreement was lower for domains that are central to behavioural inference but hard to judge from published reports, notably pseudoreplication and procedural equivalence. Pseudoreplication is a recurring concern because the effective experimental unit is often clustered, for example, within dyads, cages, or litters, yet analyses and reporting may still treat individual observations as independent. This is especially relevant for reciprocal interaction designs, where behaviour arises from a coupled pair and non-independence can inflate apparent precision and produce overconfident estimates if it is not handled appropriately in the analysis (Lazic, 2010; Lazic *et al*., 2018, 2020). Procedural equivalence is difficult to appraise retrospectively because many determinants of social behaviour depend on details that are inconsistently reported, including partner preparation and matching, handling and habituation, apparatus configuration, scoring definitions, and whether groups received the same vehicles, routes of administration, and administration to test timings apart from the intervention itself. Accordingly, disagreement for pseudoreplication and procedural equivalence likely reflects more than incomplete reporting alone. These issues are still unevenly recognised and operationalised across fields, as they are not always treated as core validity threats in routine behavioural practice, even though guidance on rigour and transparent reporting in animal research has become increasingly explicit (Lazic *et al*., 2018; Percie du Sert *et al*., 2020; Smith, 2020; Wilson *et al*., 2023).

Sex representation in the available evidence is highly imbalanced, which limits how confidently the reported NMDAR antagonist effects on social outcomes can be generalised. Sex bias is a long-standing feature of preclinical neuroscience (Beery & Zucker, 2011; Shansky & Woolley, 2016). In preclinical schizophrenia research, this imbalance is sometimes justified by concerns about variability in hormonal regulation and behaviour of females (Kokras & Dalla, 2014; Raimondi, Tripp & Ostroff, 2023), and is sometimes framed as less problematic because epidemiological evidence suggests a higher incidence of this disorder in males (McGrath *et al*., 2004; Leucht *et al*., 2025). However, the variability rationale is not well supported across many behavioural endpoints (Prendergast, Onishi & Zucker, 2014), and policy shifts have explicitly encouraged routine consideration of sex as a biological variable rather than treating male-only sampling as the default (Clayton & Collins, 2014). In this context, the predominance of male-only datasets should be treated as a constraint on external validity and on the identification of boundary conditions. A straightforward implication for this field is that future work should more routinely include both sexes and report sex-stratified results where feasible to clarify whether social phenotypes and pharmacological sensitivity are robust across sex, strengthening the generalisability of the findings.

Species coverage is a further limitation for external validity. The evidence base in this review rests largely on rodent protocols, with a smaller zebrafish subset. Although we screened records involving other species, including primates (Mao *et al*., 2008) and invertebrate models (Moulin *et al*., 2023; Dunham *et al*., 2024), these studies did not meet the inclusion criteria for controlled social interaction or social preference outcomes with extractable group-level data. As a result, the conclusions are anchored in a narrow set of species and assay traditions. A natural implication is that broader cross-species sampling, alongside deliberate introduction of heterogeneity within species and across laboratories, would help clarify which NMDAR antagonist-related social effects are robust and which are contingent on particular implementations (Richter *et al*., 2010, 2011; Voelkl *et al*., 2020; Carneiro *et al*., 2023).

A limitation of this review is that the searches were completed in 2023 in a field that continues to grow, as NMDAR antagonist models remain widely used. It is therefore likely that additional eligible studies have been published since the search date. Even so, the number of included studies was large, which strengthens the synthesis and allows a broad characterisation of the evidence base available up to that point. At the same time, inference is constrained by substantial heterogeneity in methodology and reporting, much of which could not be accounted for by the moderators that could be analysed with adequate support. In parts of the dataset, sensitivity to small study effects suggests that conventional pooled estimates may overstate effects, and risk of bias assessments were often dominated by unclear judgments because key methodological details were not reported consistently. Taken together, these limitations indicate that the pooled effects summarise a heterogeneous literature in which social behaviour is operationalised in multiple ways, rather than providing a single estimate that applies uniformly across settings.

Within these limitations, this review synthesises a large preclinical literature and summarises how NMDAR antagonism relates to social outcomes as they are most commonly assessed. By integrating findings across compounds, species, and paradigms, it provides an updated account of the consistency of reported effects and the degree of uncertainty that accompanies them. Bringing this work together also helps place individual study results in a broader context and provides a clearer basis for interpreting the wider preclinical literature on NMDAR antagonist models.

## V. CONCLUSIONS

1. Across preclinical studies, NMDAR antagonists are associated with reduced social behaviour on average, but effects are highly variable across experiments and contexts;
2. In social interaction tests, the overall evidence supports a robust reduction in reciprocal social interaction, even when accounting for small study effects;
3. In social preference paradigms, estimates also suggest reduced sociability, but bias adjustment eliminates the effect, indicating that the evidence is more sensitive to small-study effects;
4. Locomotor activity provides critical interpretive context. NMDAR antagonists increase activity during social interaction tests, highlighting the importance of separating social effects from non-social performance factors;
5. By contrast, locomotor activity measured within social preference paradigms shows no clear overall change, despite an overall reduction in social preference induced by NMDAR antagonists;
6. Clinically approved antipsychotics increase social endpoints in NMDAR antagonist-exposed animals, but parallel reductions in locomotion suggest that apparent normalisation may arise from altered activity and time allocation, which limits the extent to which these findings can be interpreted as modelling clinically meaningful improvements in social functioning;
7. The evidence base is constrained by substantial heterogeneity and by incomplete reporting of core design and analysis safeguards, including randomisation, blinding, and unit of analysis decisions, as well as marked sex imbalance;
8. Future studies will be most informative if they strengthen transparency and methodological rigour, predefine primary social outcomes, routinely report within-task activity measures alongside social endpoints, and use designs and analyses that appropriately account for non-independent observations.

## ACKNOWLEDGEMENTS

We thank Dr. Dirce Maria Santin (orcid.org/0000-0003-1721-5115), Librarian at Instituto de Ciências Básicas da Saúde (UFRGS), for her help in reviewing the search strategies. We also thank the BRISA team for helpful comments and suggestions throughout the work. This study was supported by grants from Coordenação de Aperfeiçoamento de Pessoal de Nível Superior (CAPES), Conselho Nacional de Desenvolvimento Científico e Tecnológico (CNPQ, grant number 401800/2025-3), Pró-Reitoria de Pesquisa (UFRGS), and Fundação de Amparo à Pesquisa do Estado do Rio Grande do Sul (FAPERGS, grant number 24/2551-0001330-0). The funders had no role in the study design, data collection, data analysis, data interpretation, manuscript preparation, or the decision to submit the article for publication. The authors declare no competing interests.

Assistance from ChatGPT 5.2 (OpenAI, San Francisco, CA, USA) was used to support the optimisation of text and formatting throughout the manuscript. All outputs were critically reviewed, edited, and validated by the authors, who take full responsibility for the content and interpretation of the work.

## VI. AUTHOR CONTRIBUTIONS

**Conceptualization:** MG-L, MBA, MA, BDA, LMB, TCC, DVM, AP-A, DAR, DJS, APH; **Data curation:** MG-L, APH; **Formal Analysis:** MG-L; **Funding acquisition:** APH; **Investigation:** MG-L, MBA, MA, BDA, LMB, TCC, DVM, AP-A, DAR, DJS, APH; **Methodology:** MG-L, MBA, MA, BDA, LMB, TCC, DVM, AP-A, DAR, DJS, APH; **Project administration:** MG-L, APH; **Resources:** APH; **Software:** MG-L; **Supervision:** APH; **Validation:** MG-L; **Visualization:** MG-L; **Writing – original draft:** MG-L; **Writing – review & editing:** MBA, MA, BDA, LMB, TCC, DVM, AP-A, DAR, DJS, APH.

## VII. DATA ACCESSIBILITY

Data and code supporting the findings of this study are openly available via the Open Science Framework <u>osf.io/98fau</u> (Gallas-Lopes *et al*., 2023a) and GitHub <u>github.com/matheusglls/NMDA-sb</u> (Gallas-Lopes, 2026).

