## Supplementary Material for "Effects of NMDA antagonists on social behaviour: a systematic review and meta-analysis of preclinical studies"

[1]Brazilian Reproducibility Initiative in preclinical Systematic review and meta-Analysis (BRISA), Rio de Janeiro, Brazil.

[2]Universidade Federal do Rio Grande do Sul (UFRGS), Porto Alegre, Brazil.

[3]Universidade Federal do Rio de Janeiro (UFRJ), Rio de Janeiro, Brazil.

[4]Hospital de Clínicas de Porto Alegre (HCPA), Porto Alegre, Brazil.

[5]Universidade Federal de Goiás (UFG), Goiânia, Brazil.

*Corresponding author: Ana Paula Herrmann. Departamento de Farmacologia, Instituto de Ciências Básicas da Saúde (ICBS), Universidade Federal do Rio Grande do Sul (UFRGS), Rua Ramiro Barcelos 2600/430, Porto Alegre, RS, 90610-264, Brazil.. Telephone: +55 51 33082066

**Figure S1**

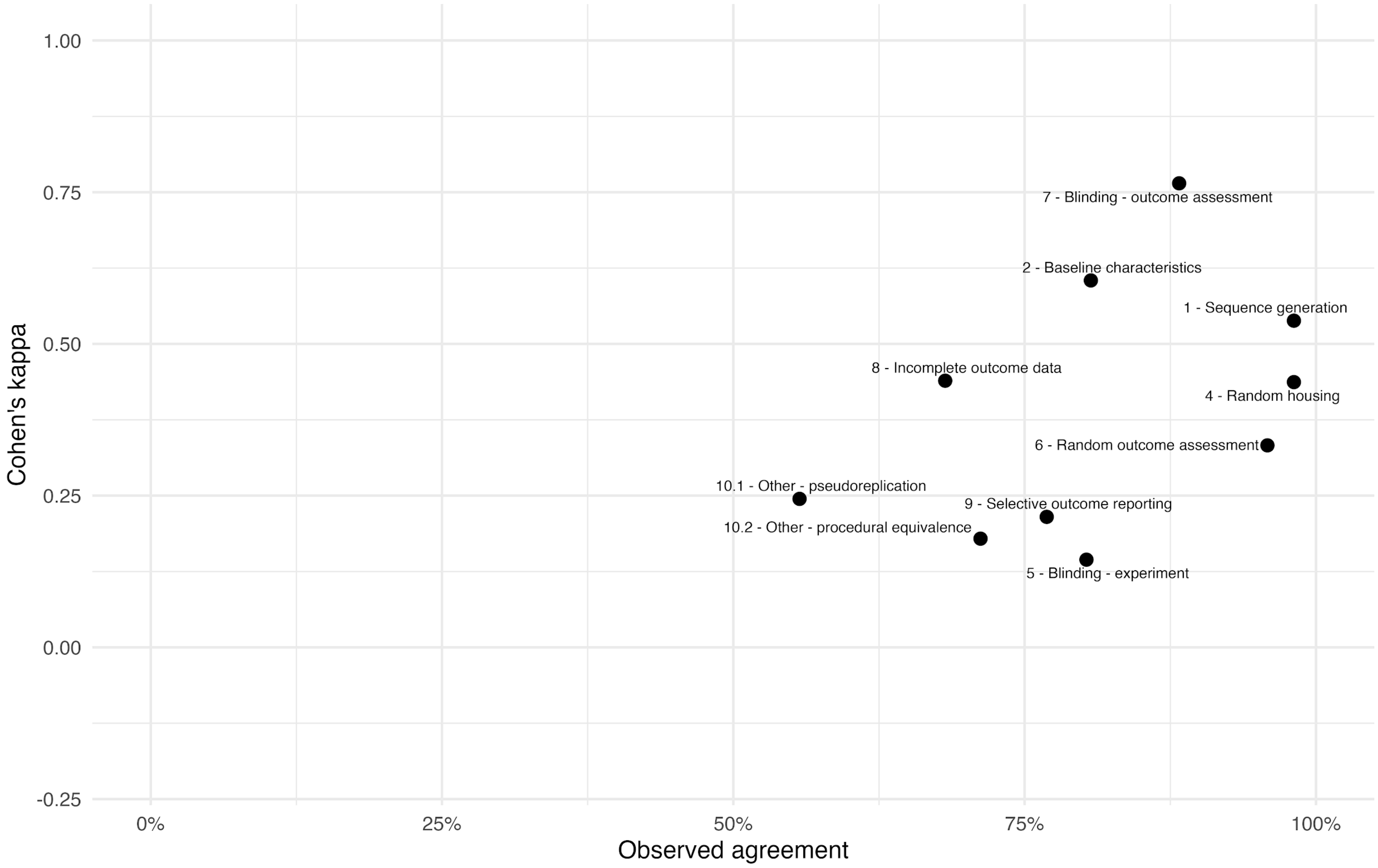


**Figure S1. Observed agreement plotted against Cohen's kappa for each Risk of Bias domain.** Each point represents one domain from the risk of bias assessment based on independent ratings of two reviewers. The x-axis shows the percentage of identical judgments between reviewers (observed agreement), whereas the y-axis shows Cohen's kappa, which corrects for agreement expected by chance.

**Figure S2**

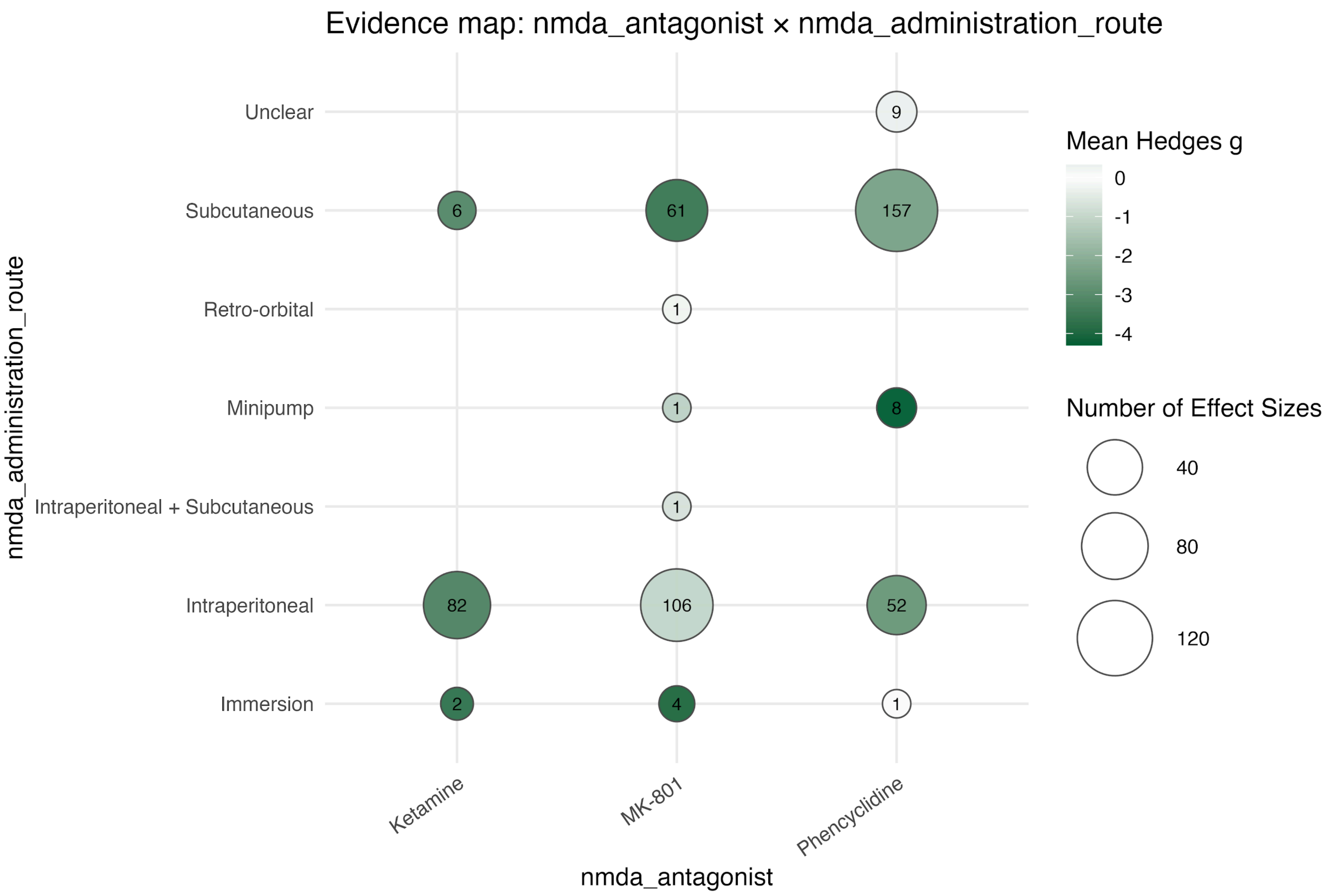


**Figure S2. Evidence map of effect sizes for social interaction by NMDA antagonist and route of administration.** Circle size represents the number of effect sizes, and colour intensity indicates the mean Hedges' g.

**Figure S3**

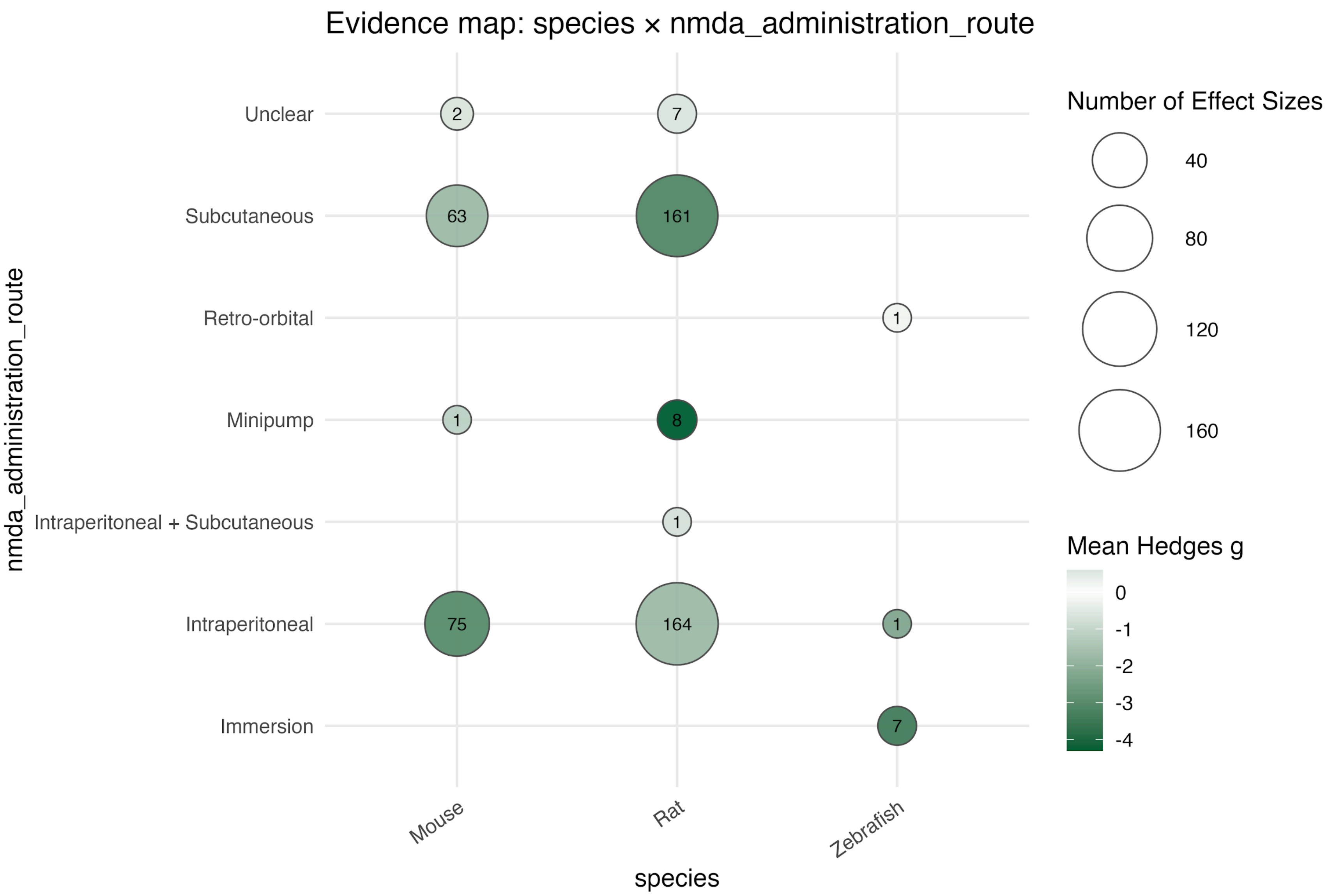


**Figure S3. Evidence map of effect sizes for social interaction by species and route of administration.** Circle size represents the number of effect sizes, and colour intensity indicates the mean Hedges' g.

**Figure S4**

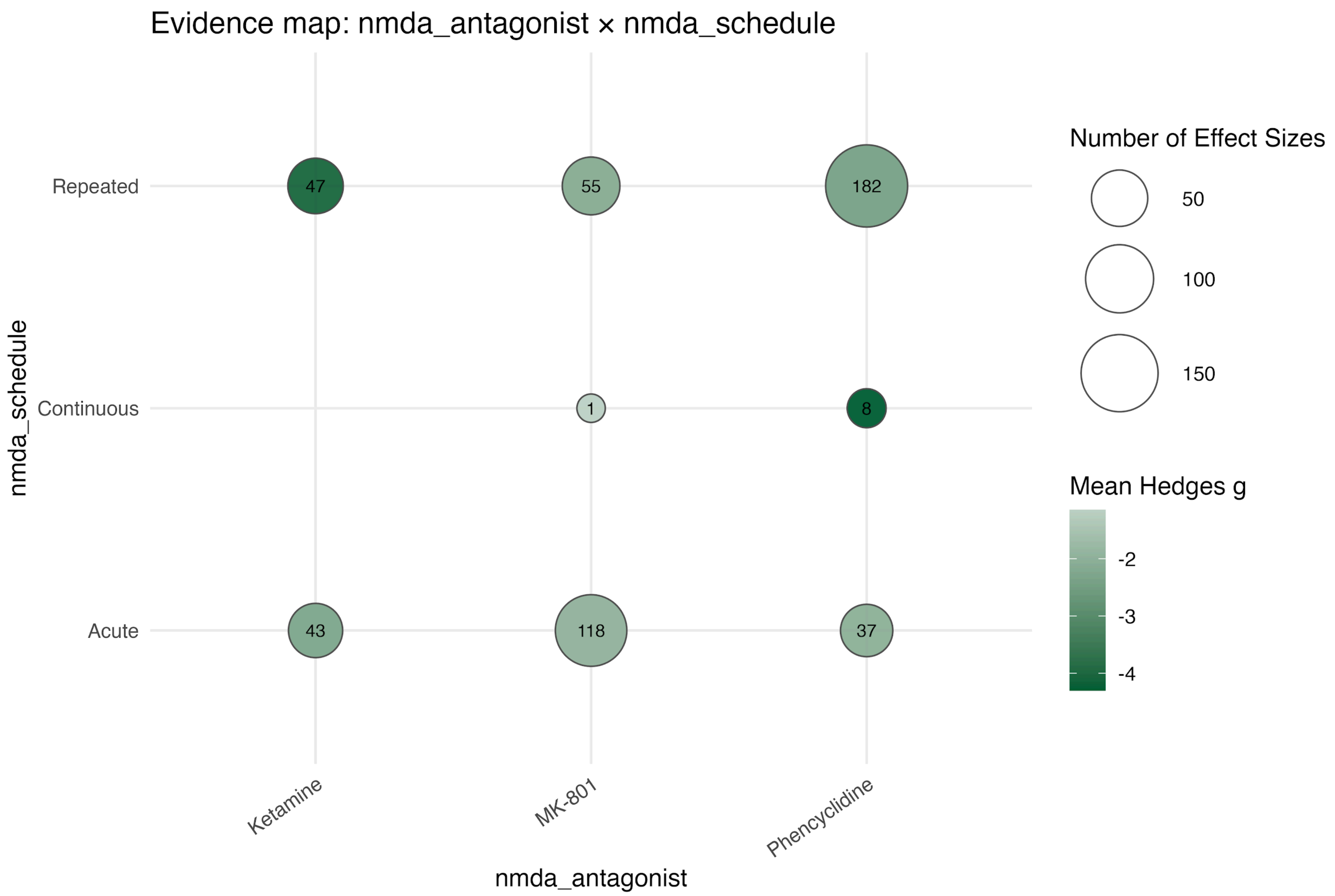


**Figure S4. Evidence map of effect sizes for social interaction by NMDA antagonist and schedule of administration.** Circle size represents the number of effect sizes, and colour intensity indicates the mean Hedges' g.

**Figure S5**

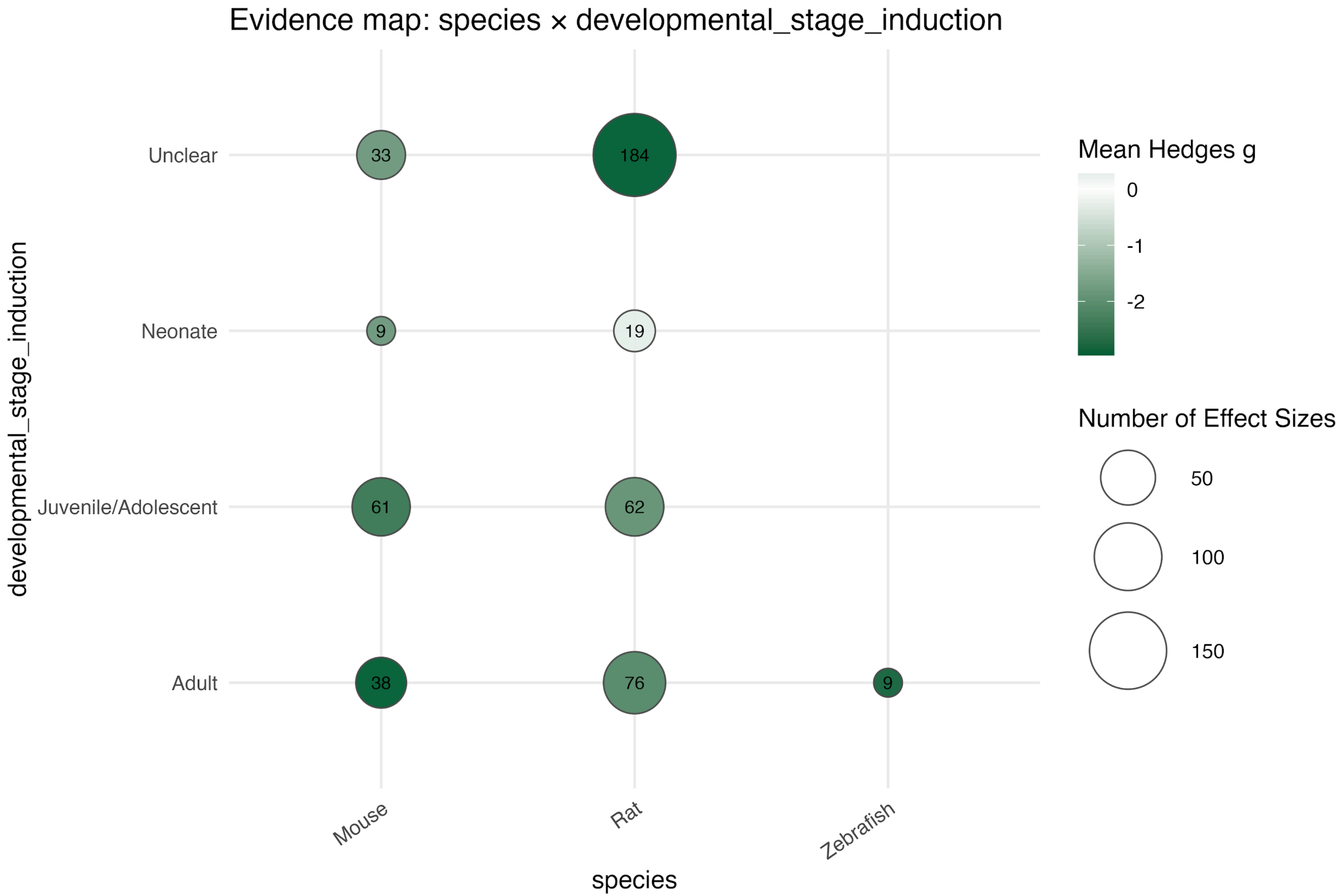


**Figure S5. Evidence map of effect sizes for social interaction by species and developmental stage at induction.** Circle size represents the number of effect sizes, and colour intensity indicates the mean Hedges' g.

**Figure S6**

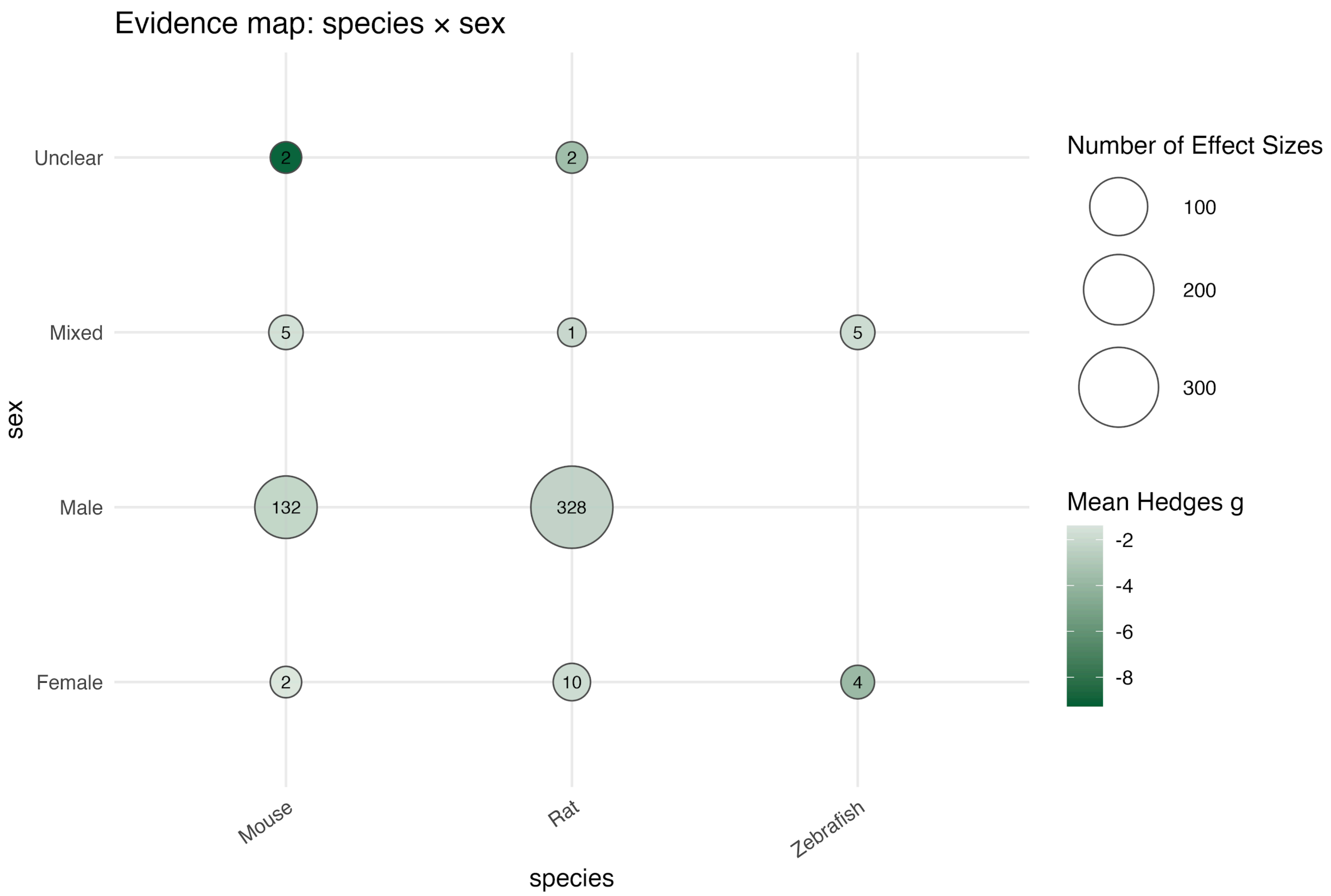


**Figure S6. Evidence map of effect sizes for social interaction by species and sex.** Circle size represents the number of effect sizes, and colour intensity indicates the mean Hedges' g.

**Figure S7**

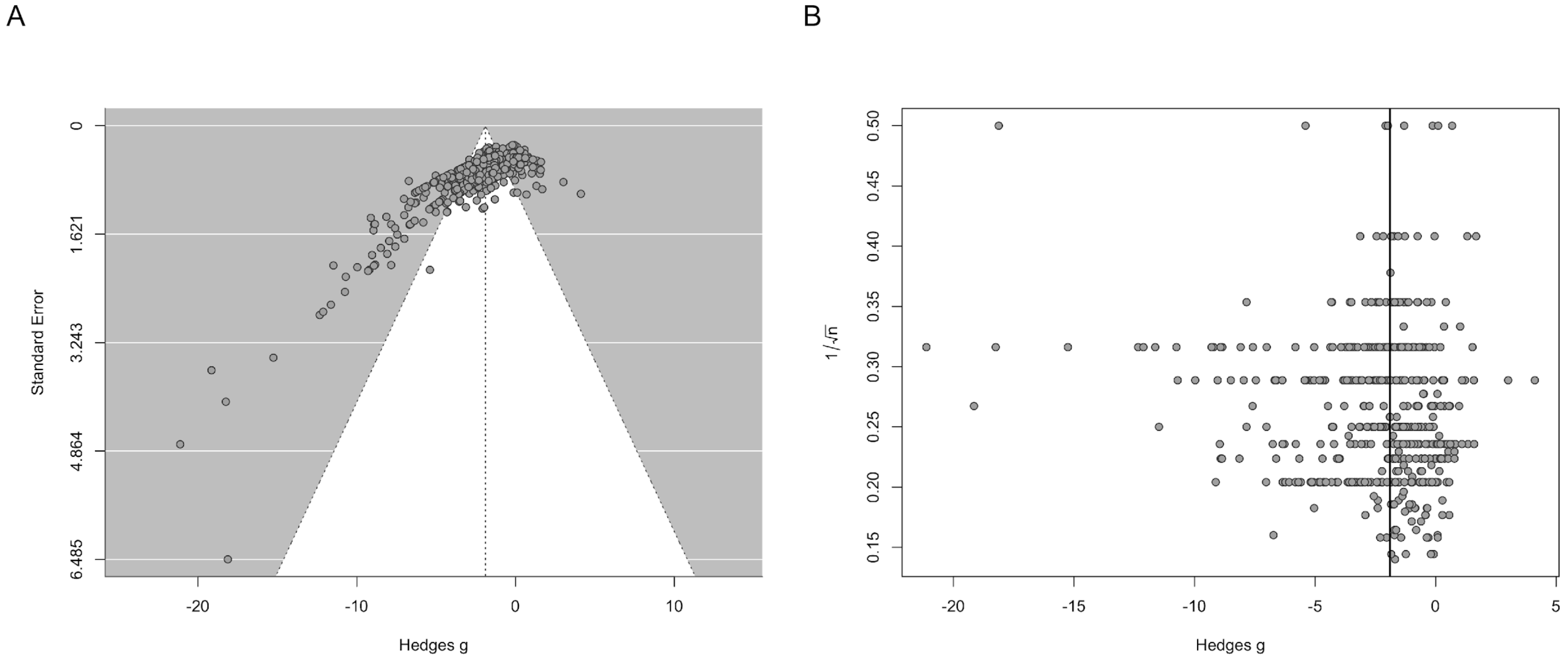


**Figure S7. Funnel plots of effect sizes for social interaction.** (A) Funnel plot showing Hedges' g plotted against the standard error. (B) Funnel plot showing Hedges' g plotted against the inverse square root of total sample size as an alternative measure of study precision. Each point corresponds to one effect size.

**Figure S8**

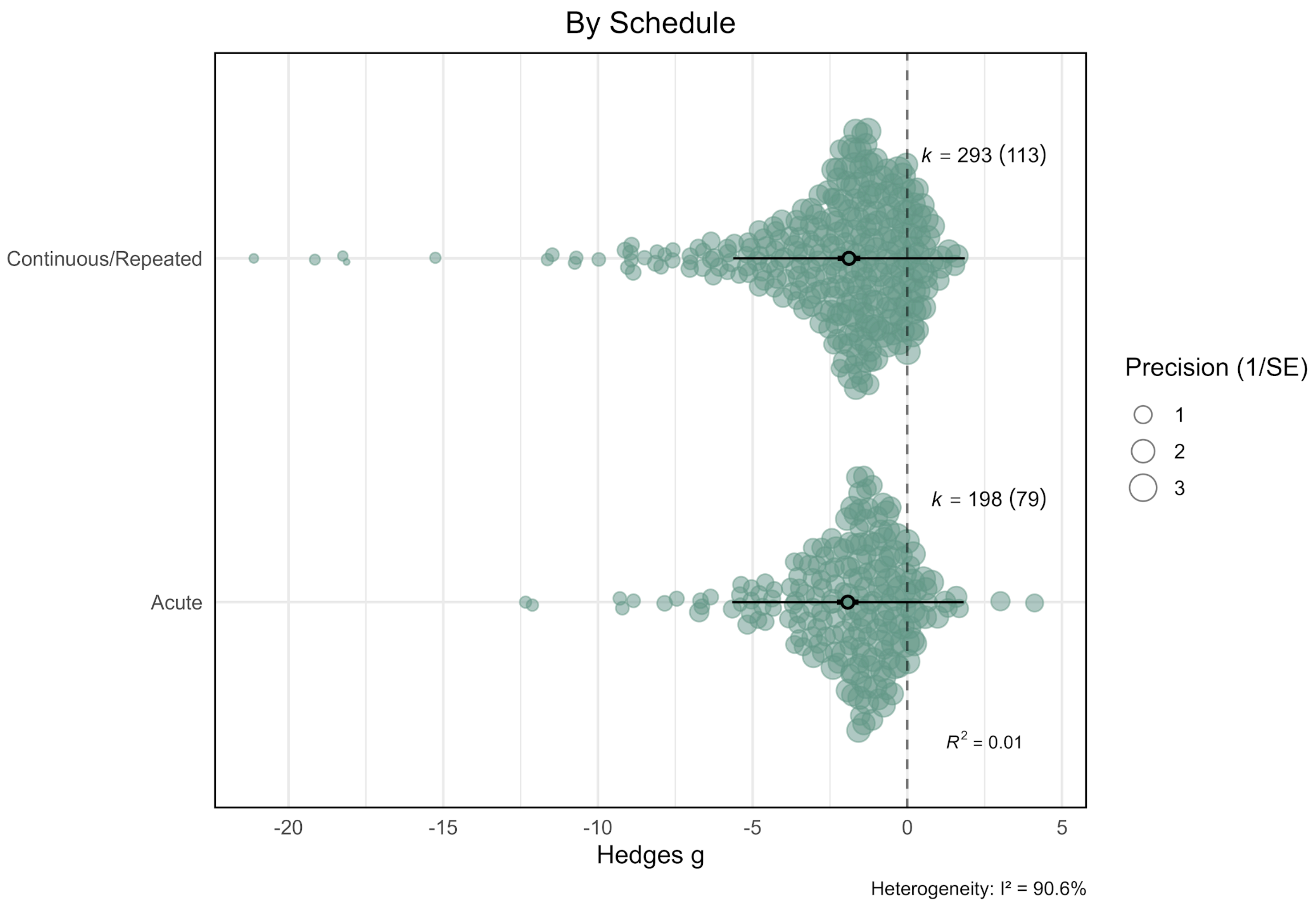


**Figure S8. Effects of NMDA receptor antagonists on social interaction stratified by administration schedule.** Orchard plot showing pooled effects for acute and continuous/repeated schedules. Filled circles represent individual effect sizes scaled by study precision, and open circles indicate pooled point estimates with 95% confidence intervals (thick horizontal lines) and prediction intervals (thin horizontal lines). k denotes the number of effect sizes, with the number of independent studies shown in parentheses.

**Figure S9**

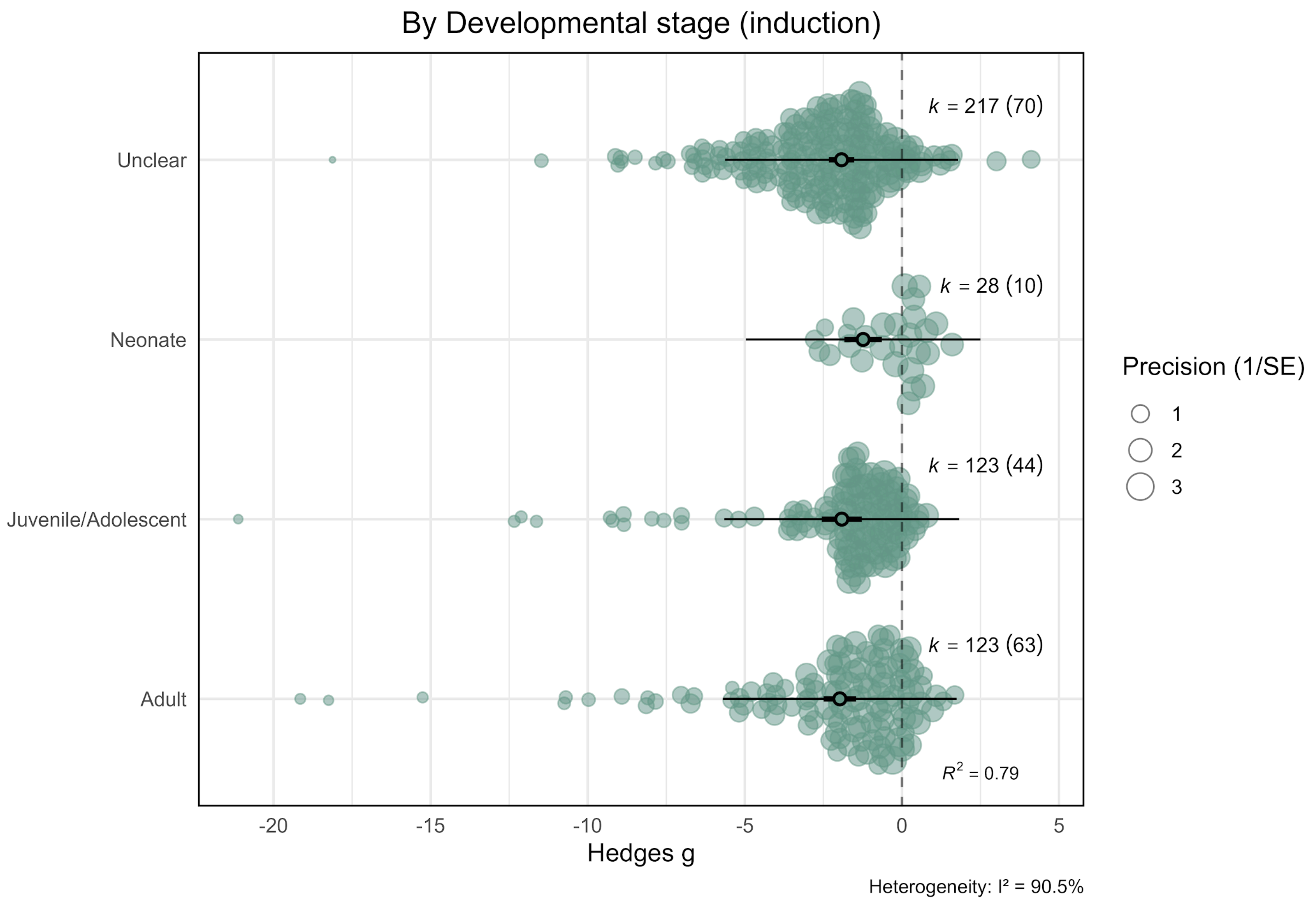


**Figure S9. Effects of NMDA receptor antagonists on social interaction stratified by developmental stage at induction.** Orchard plot showing pooled effects for adult, juvenile/adolescent, neonatal, and studies with unclear developmental stage at induction. Filled circles represent individual effect sizes scaled by study precision, and open circles indicate pooled point estimates with 95% confidence intervals (thick horizontal lines) and prediction intervals (thin horizontal lines). k denotes the number of effect sizes, with the number of independent studies shown in parentheses.

**Figure S10**

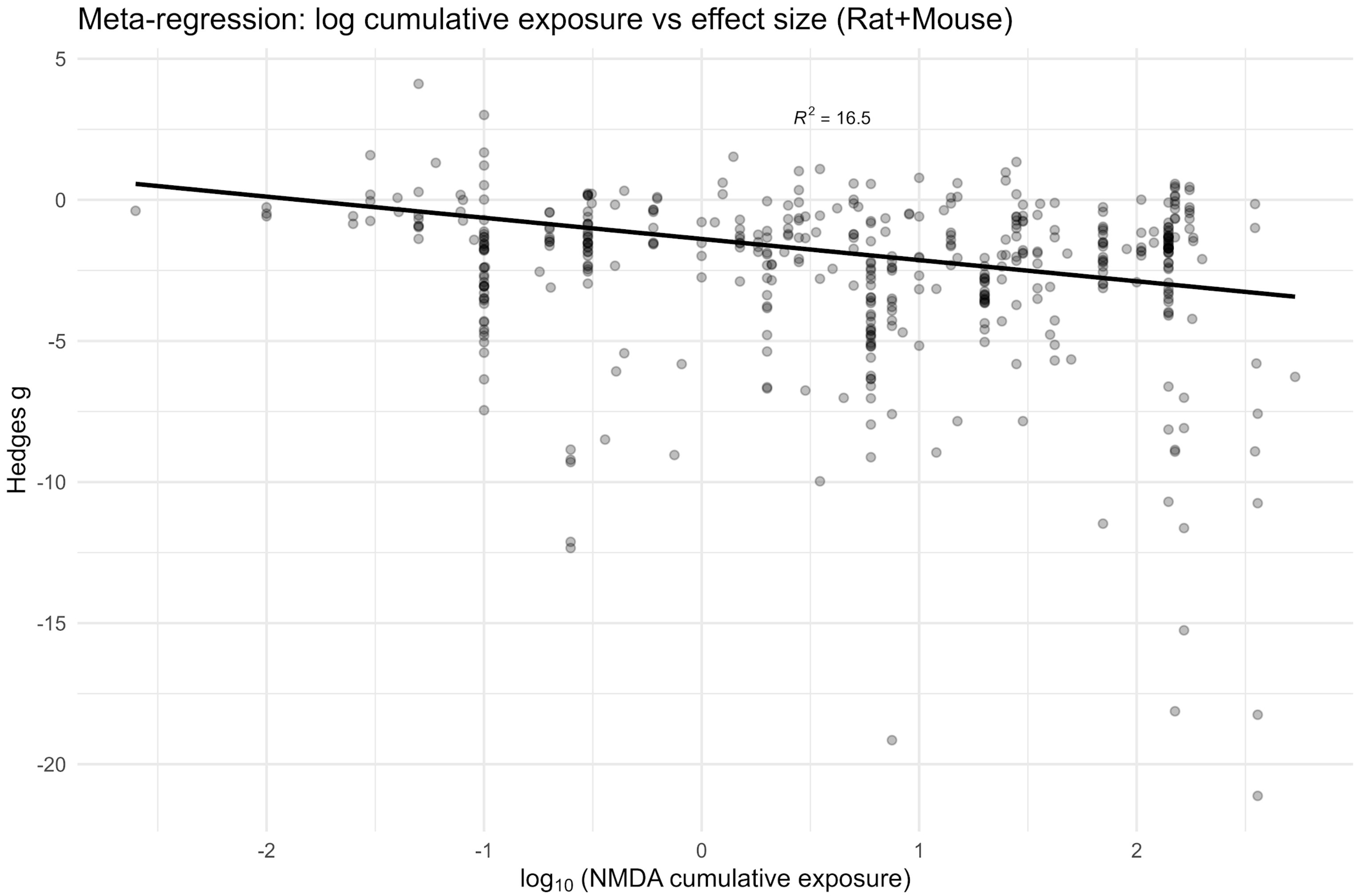


**Figure S10. Meta-regression of log-transformed cumulative NMDA antagonist exposure and effect sizes for social interaction.** The plot shows Hedges' g as a function of $\log_{10}$-transformed cumulative NMDA antagonist exposure, including only studies conducted in rats and mice. Cumulative exposure was calculated, where data were available, as the product of dose, intervention frequency, and duration of exposure. Each point represents one effect size, and the solid line represents the fitted meta-regression slope from a multivariate random-effects model.

**Figure S11**

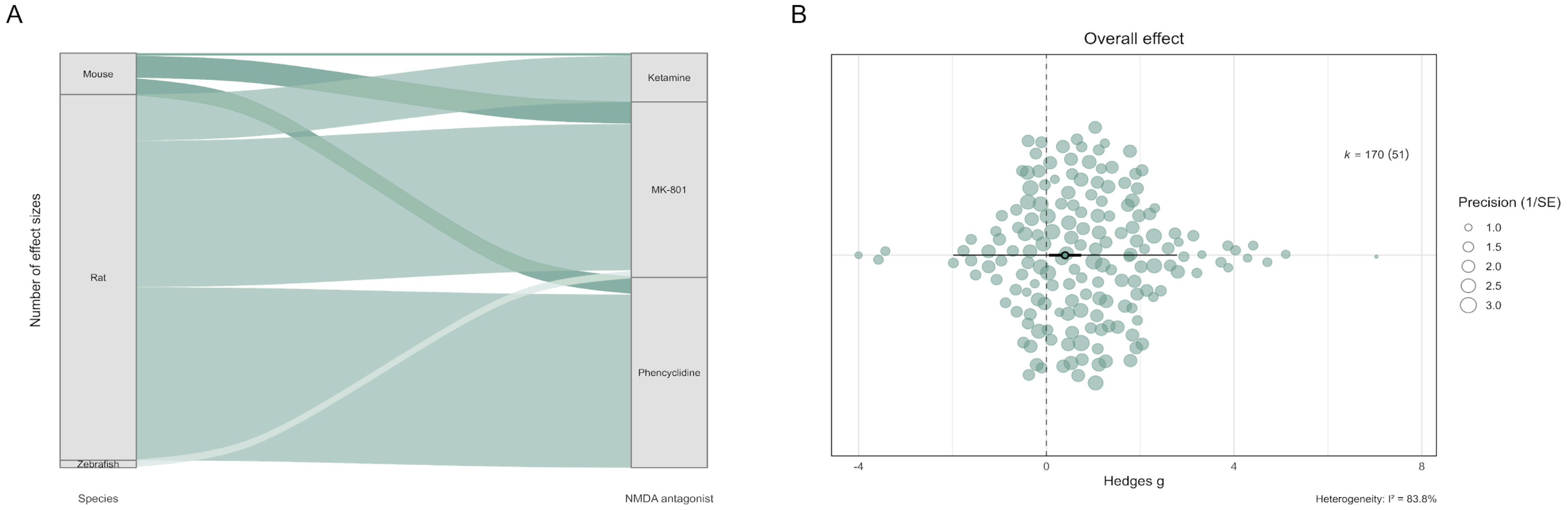


**Figure S11. Effects of NMDA receptor antagonists on locomotor activity in experiments measuring social interaction.** (A) Alluvial plot showing the distribution of effect sizes across species and NMDA receptor antagonists for studies in which locomotor activity was assessed alongside social interaction. Flow width represents the number of effect sizes contributing to each category. (B) Orchard plot of the overall pooled effect of NMDA receptor antagonists on locomotor activity. Filled circles represent individual effect sizes scaled by study precision, and the open circle indicates the pooled point estimate with 95% confidence intervals (thick horizontal lines) and prediction interval (thin horizontal lines). k denotes the number of effect sizes, with the number of independent studies shown in parentheses.

**Figure S12**

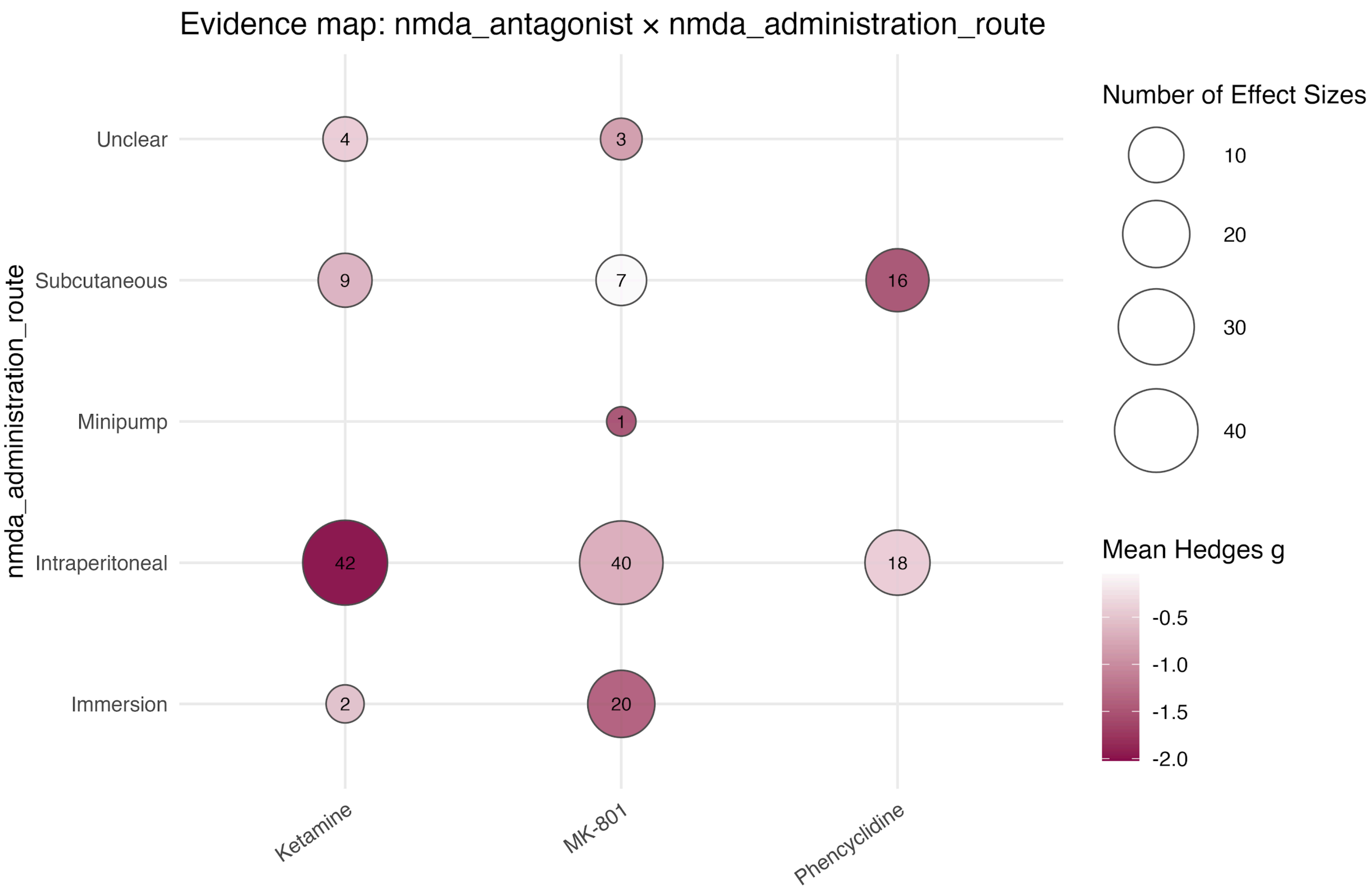


**Figure S12. Evidence map of effect sizes for social preference by NMDA antagonist and route of administration.** Circle size represents the number of effect sizes, and colour intensity indicates the mean Hedges' g.

**Figure S13**

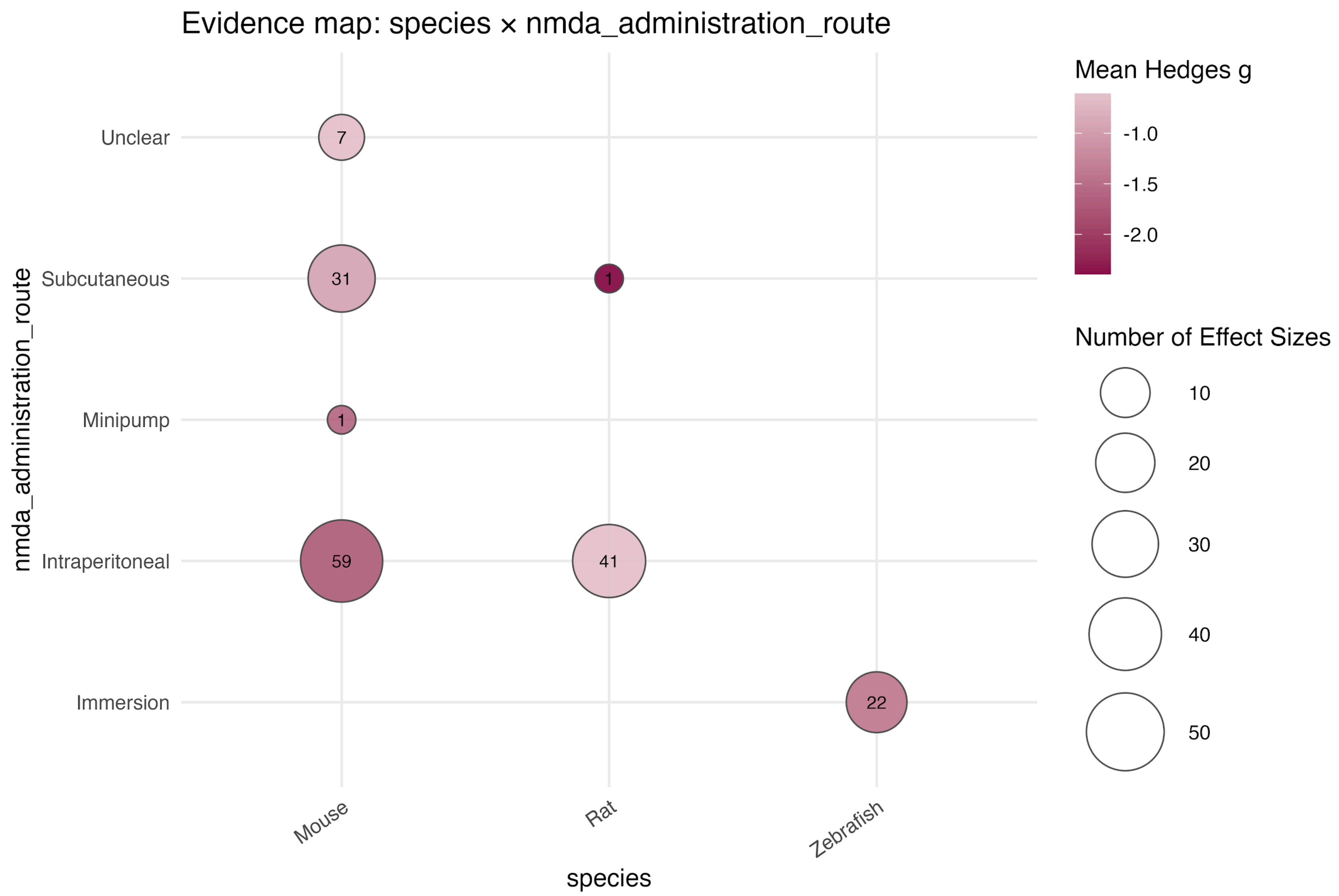


**Figure S13. Evidence map of effect sizes for social preference by species and route of administration.** Circle size represents the number of effect sizes, and colour intensity indicates the mean Hedges' g.

**Figure S14**

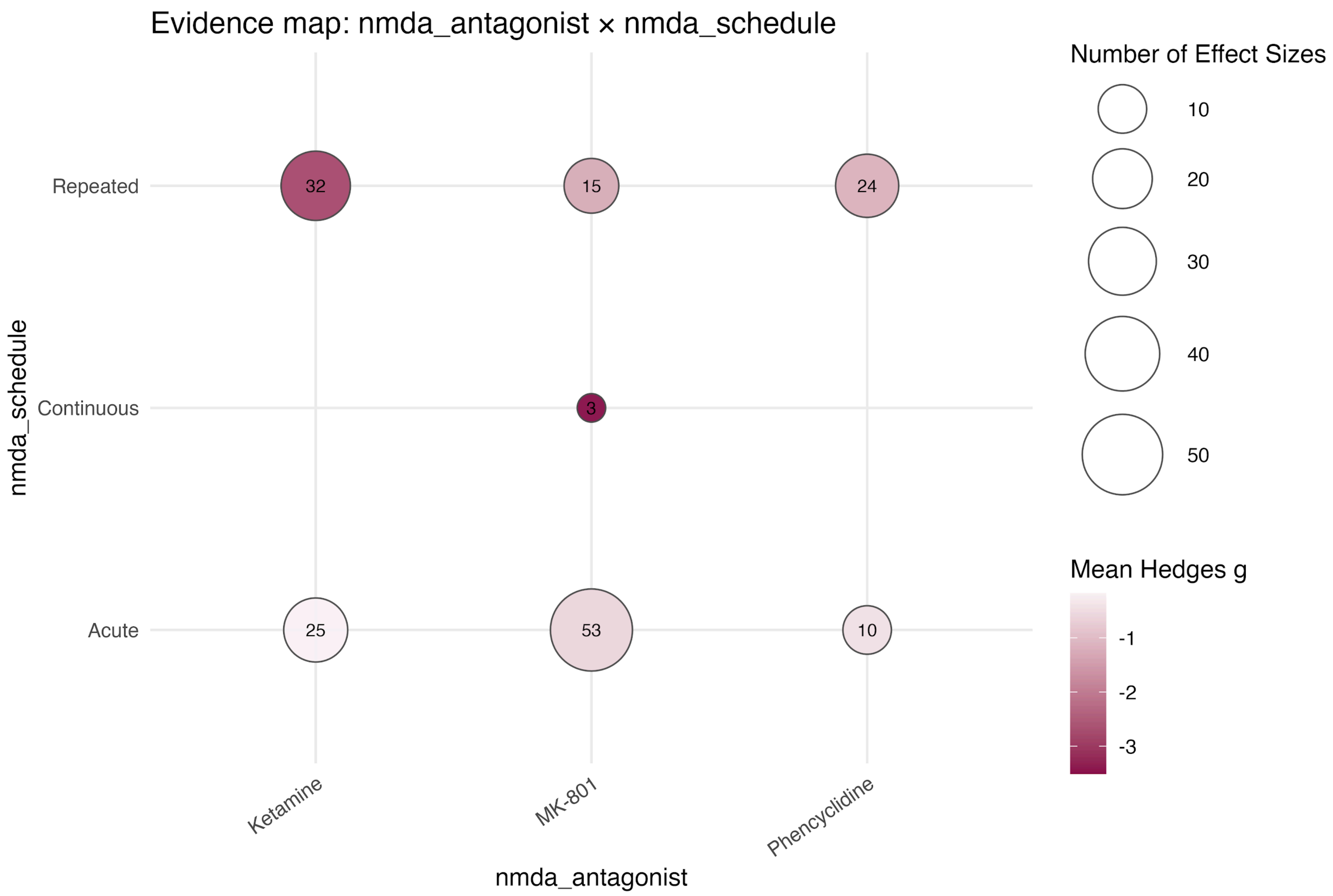


**Figure S14. Evidence map of effect sizes for social preference by NMDA antagonist and schedule of administration.** Circle size represents the number of effect sizes, and colour intensity indicates the mean Hedges' g.

**Figure S15**

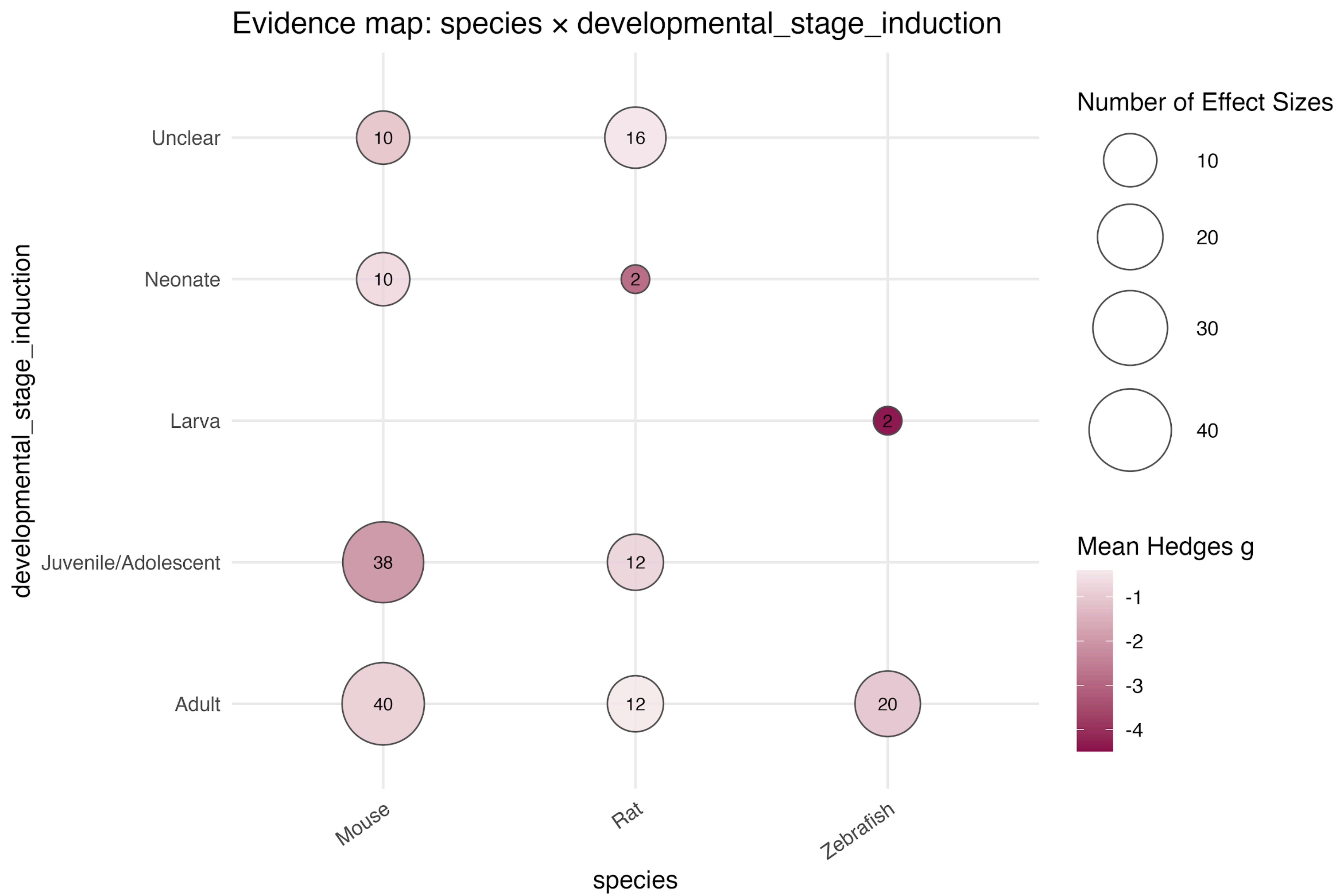


**Figure S15. Evidence map of effect sizes for social preference by species and developmental stage at induction.** Circle size represents the number of effect sizes, and colour intensity indicates the mean Hedges' g.

**Figure S16**

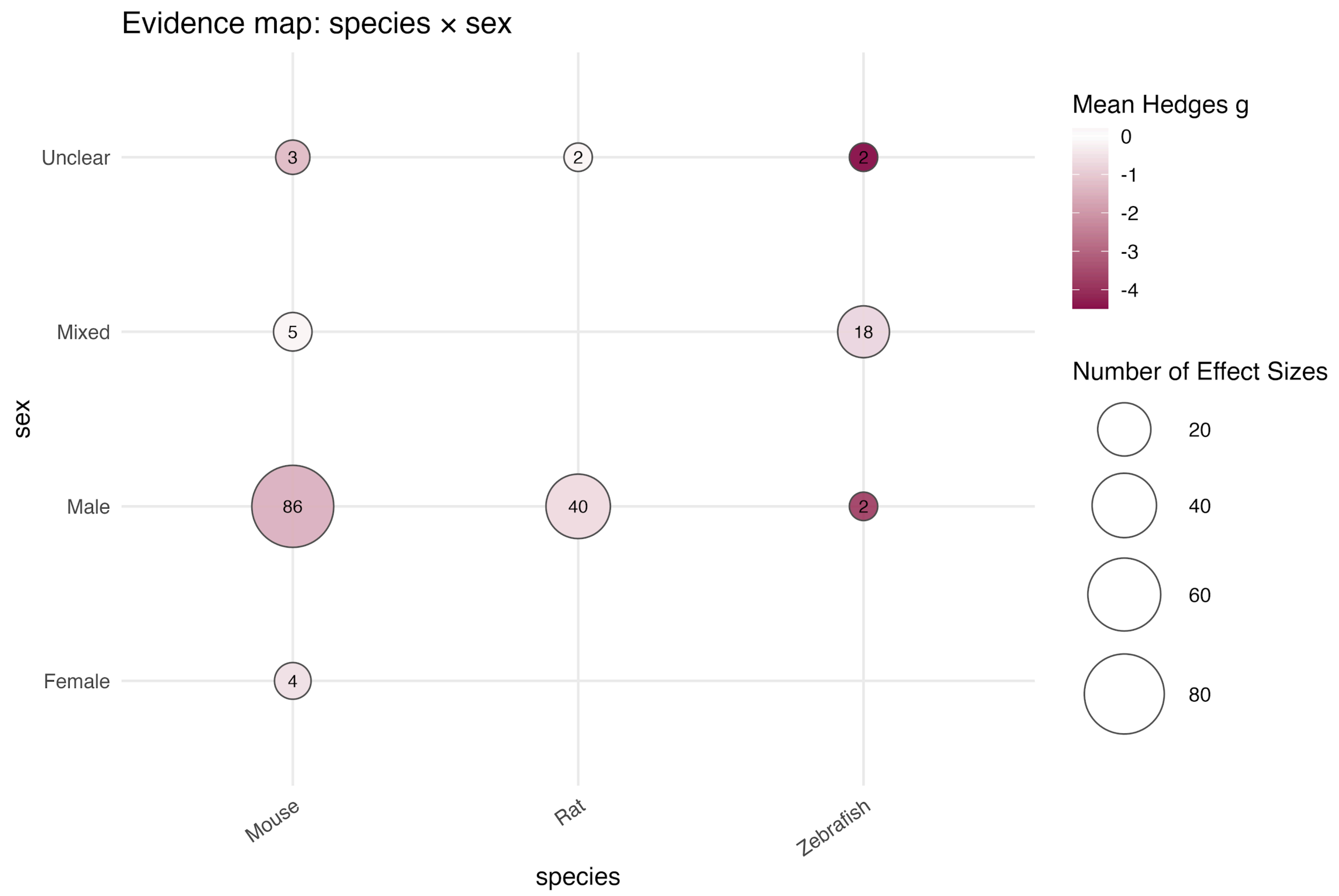


**Figure S16. Evidence map of effect sizes for social preference by species and sex.** Circle size represents the number of effect sizes, and colour intensity indicates the mean Hedges' g.

**Figure S17**

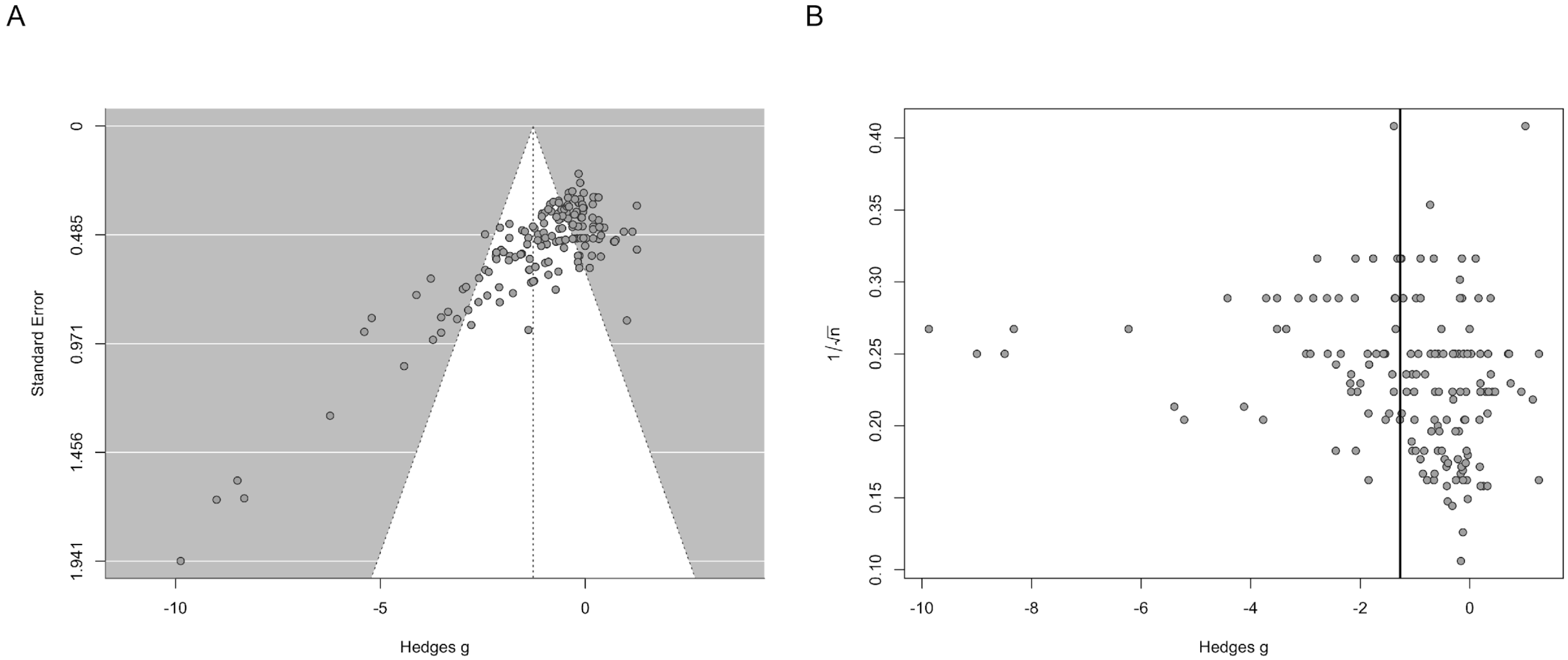


**Figure S17. Funnel plots of effect sizes for social preference.** (A) Funnel plot showing Hedges' g plotted against the standard error. (B) Funnel plot showing Hedges' g plotted against the inverse square root of total sample size as an alternative measure of study precision. Each point corresponds to one effect size.

**Figure S18**

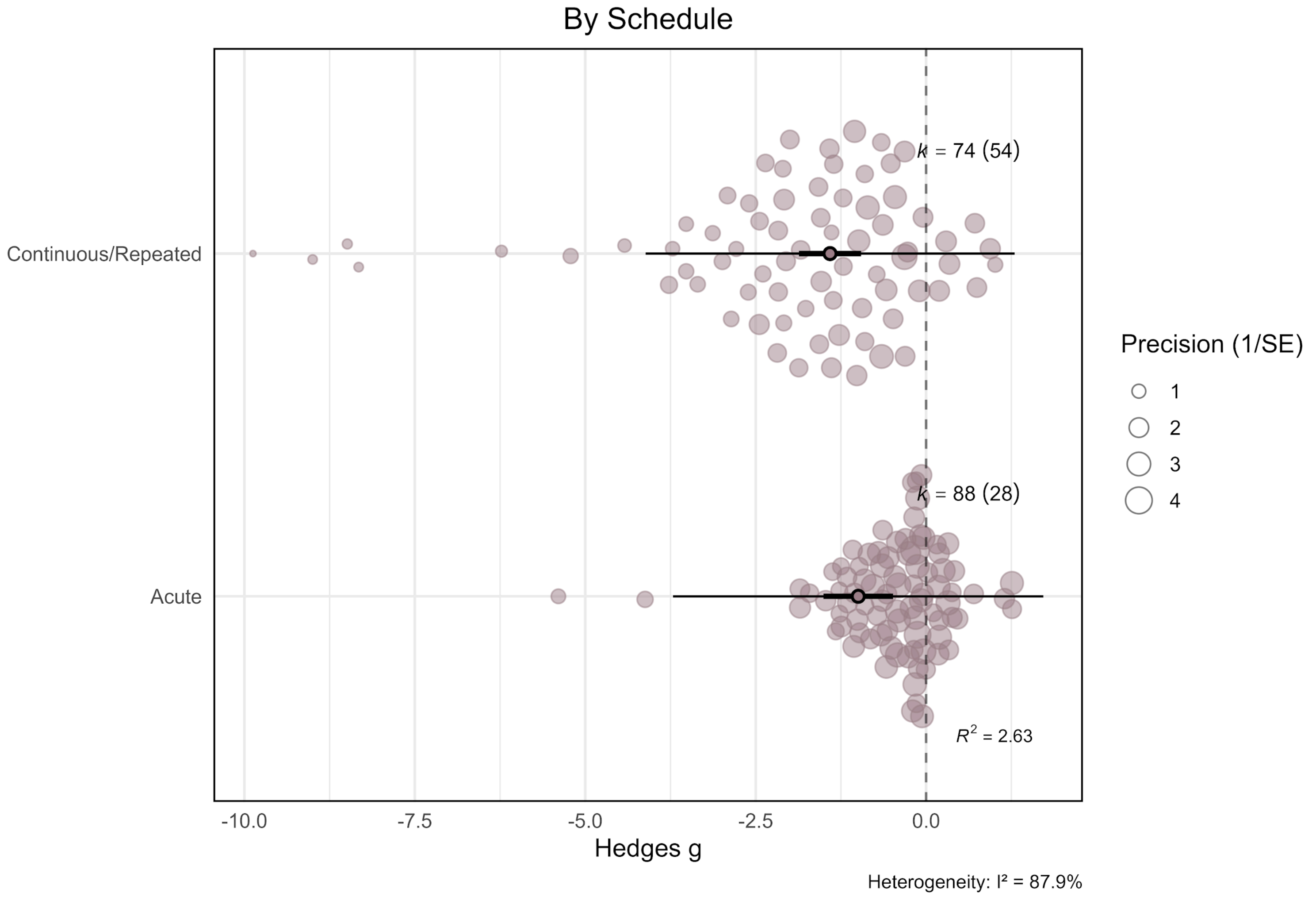


**Figure S18. Effects of NMDA receptor antagonists on social preference stratified by administration schedule.** Orchard plot showing pooled effects for acute and continuous/repeated schedules. Filled circles represent individual effect sizes scaled by study precision, and open circles indicate pooled point estimates with 95% confidence intervals (thick horizontal lines) and prediction intervals (thin horizontal lines). k denotes the number of effect sizes, with the number of independent studies shown in parentheses.

**Figure S19**

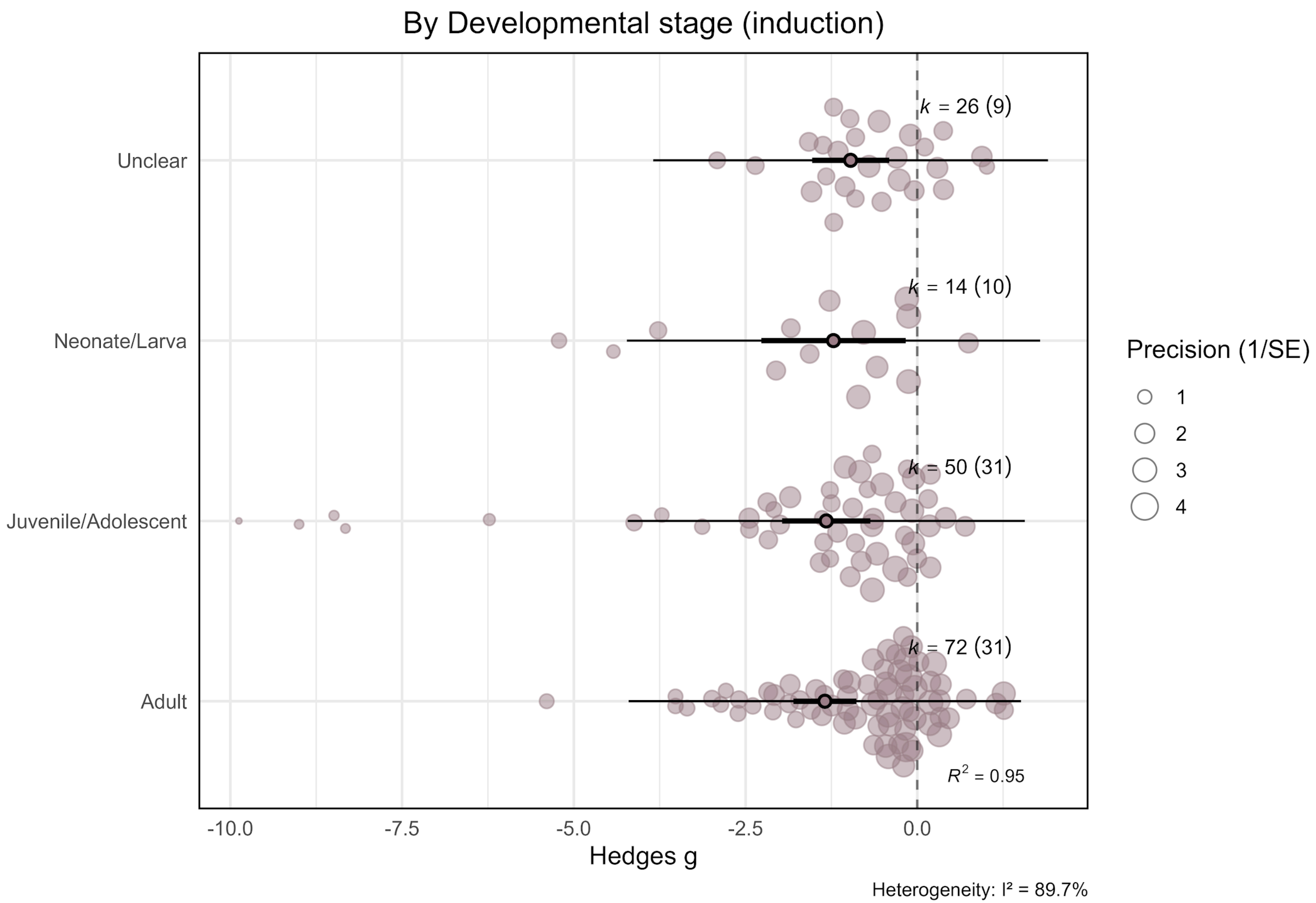


**Figure S19. Effects of NMDA receptor antagonists on social preference stratified by developmental stage at induction.** Orchard plot showing pooled effects for adult, juvenile/adolescent, neonatal/larval, and studies with unclear developmental stage at induction. Filled circles represent individual effect sizes scaled by study precision, and open circles indicate pooled point estimates with 95% confidence intervals (thick horizontal lines) and prediction intervals (thin horizontal lines). k denotes the number of effect sizes, with the number of independent studies shown in parentheses.

**Figure S20**

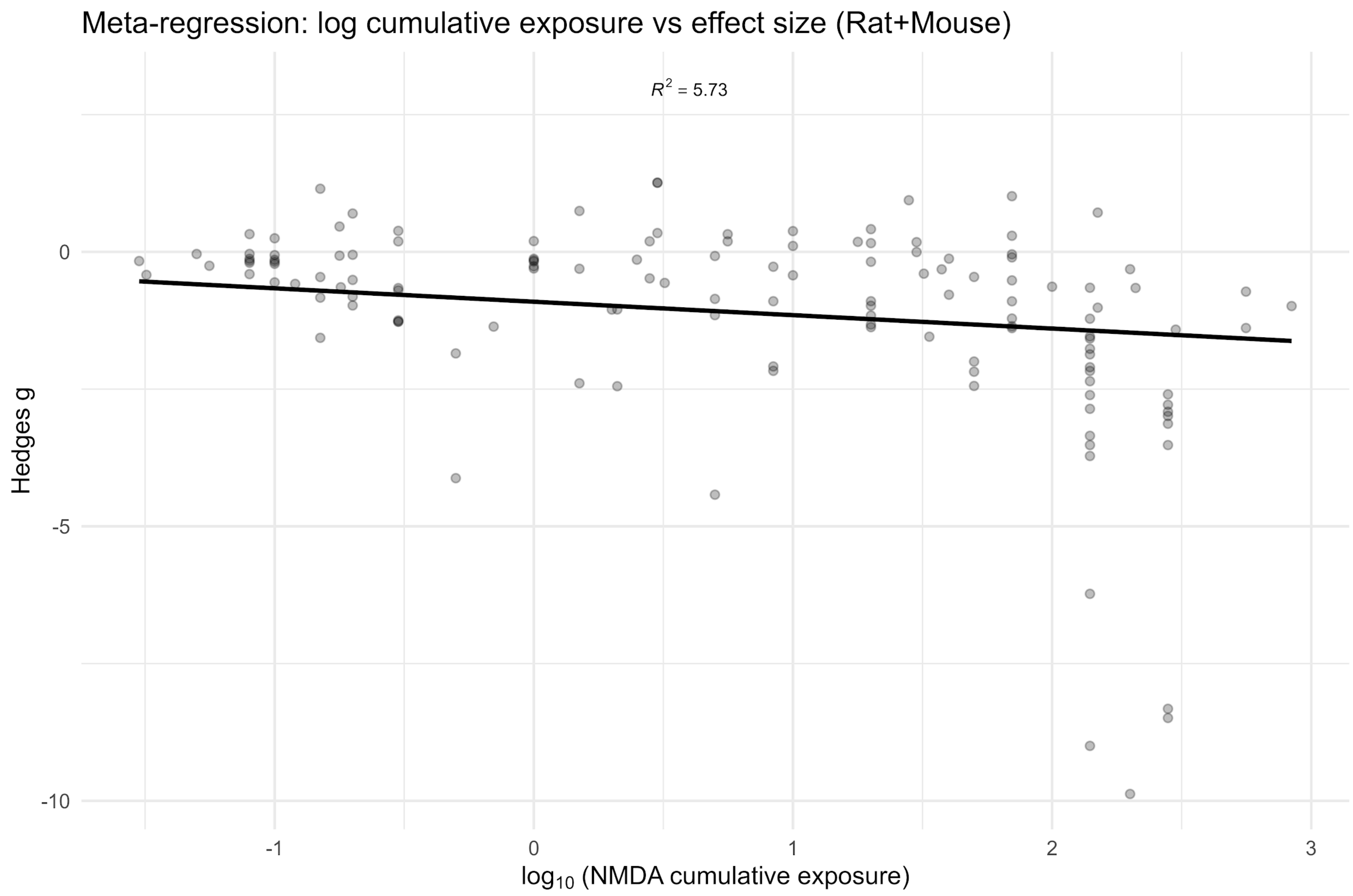


**Figure S20. Meta-regression of log-transformed cumulative NMDA antagonist exposure and effect sizes for social preference.** The plot shows Hedges' g as a function of $\log_{10}$-transformed cumulative NMDA antagonist exposure, including only studies conducted in rats and mice. Cumulative exposure was calculated, where data were available, as the product of dose, intervention frequency, and duration of exposure. Each point represents one effect size, and the solid line represents the fitted meta-regression slope from a multivariate random-effects model.

**Figure S21**

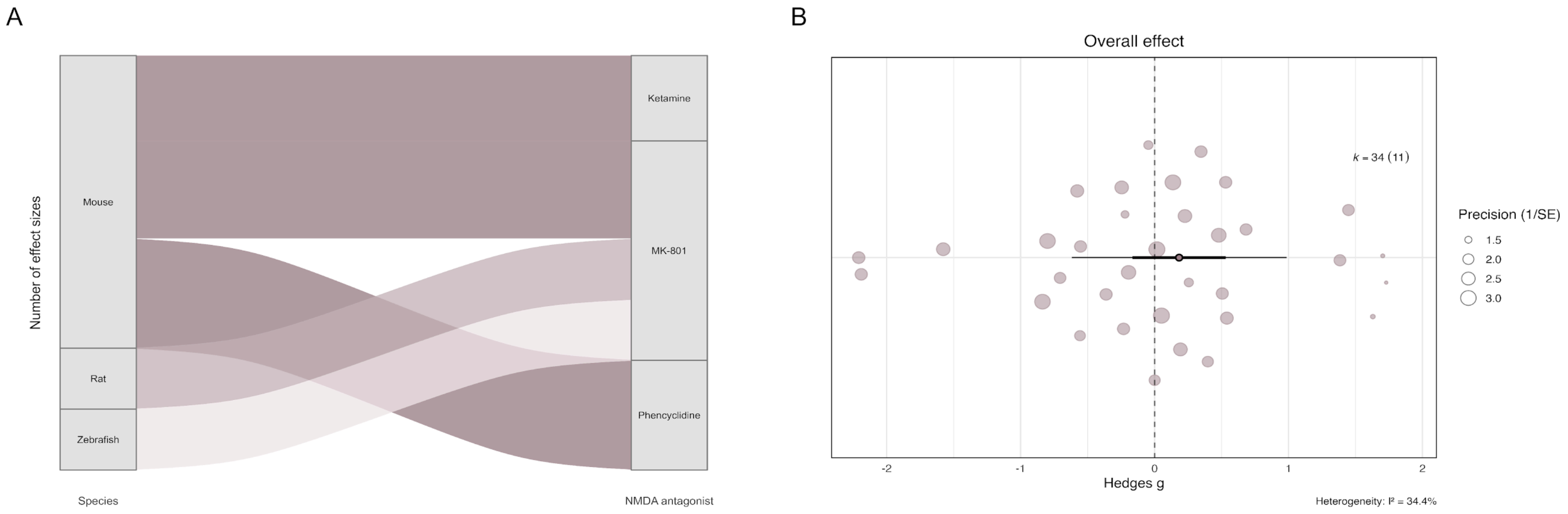


**Figure S21. Effects of NMDA receptor antagonists on locomotor activity in experiments measuring social preference.** (A) Alluvial plot showing the distribution of effect sizes across species and NMDA receptor antagonists for studies in which locomotor activity was assessed alongside social preference. Flow width represents the number of effect sizes contributing to each category. (B) Orchard plot of the overall pooled effect of NMDA receptor antagonists on locomotor activity. Filled circles represent individual effect sizes scaled by study precision, and the open circle indicates the pooled point estimate with 95% confidence intervals (thick horizontal lines) and prediction interval (thin horizontal lines). k denotes the number of effect sizes, with the number of independent studies shown in parentheses.

**Figure S22**

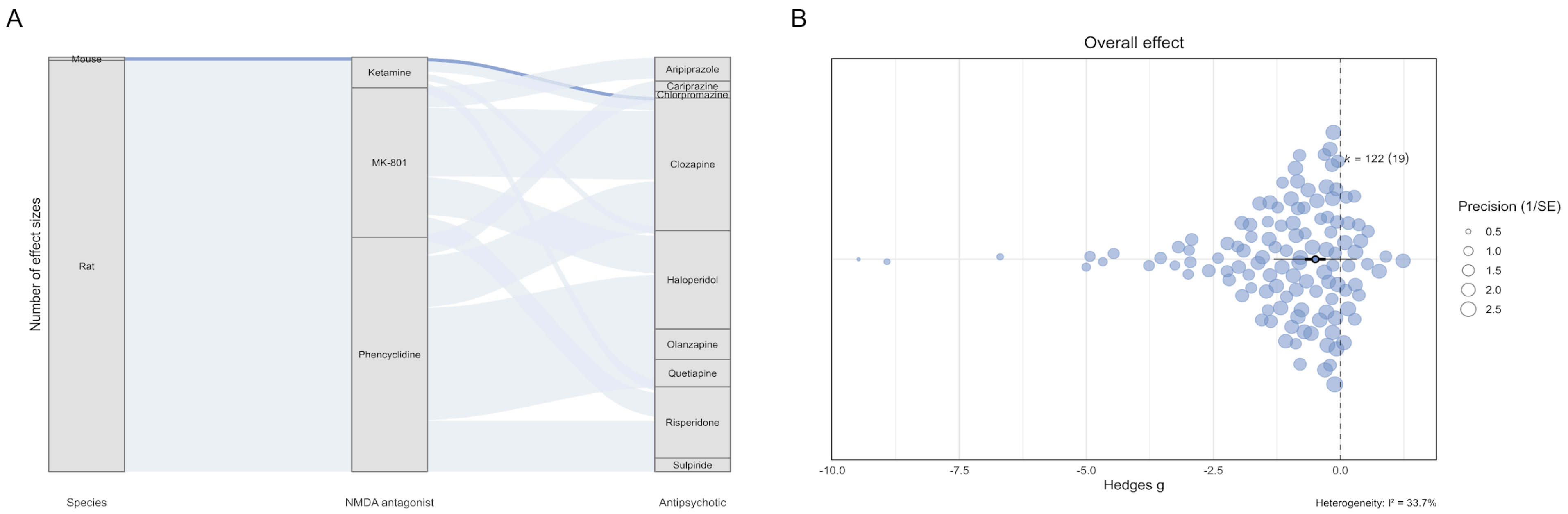


**Figure S22. Effects of antipsychotics on NMDA receptor antagonist–induced locomotor alterations in social interaction experiments.** (A) Alluvial plot showing the distribution of effect sizes across species, NMDA receptor antagonists, and antipsychotics for experiments in which locomotor activity was assessed alongside social interaction. Flow width represents the number of effect sizes contributing to each category. (B) Orchard plot of the overall pooled effect of antipsychotics on NMDA antagonist–induced locomotor alterations. Filled circles represent individual effect sizes scaled by study precision, and the open circle indicates the pooled point estimate with 95% confidence intervals (thick horizontal lines) and prediction interval (thin horizontal lines). k denotes the number of effect sizes, with the number of independent studies shown in parentheses.
